# Dysferlin is a novel regulator of COMP-positive matrifibrocytes in heart failure

**DOI:** 10.64898/2026.08.18.745492

**Authors:** Ievgeniia Kocherova, Muriel Giger, Andrea Laimbacher, Lukas Minder, Daria Nurzynska, Franca Di Meglio, Gino A Bonazza, Elena Pachera, Filip Rolski, Michał Mączewski, Przemyslaw Leszek, Michele Visentin, Oliver Distler, Przemysław Błyszczuk, Gabriela Kania

**Affiliations:** Center of Experimental Rheumatology, Department of Rheumatology, University Hospital Zurich, University of Zurich, Zurich, Switzerland; Department of Medicine, Surgery and Dentistry, University of Salerno, Baronissi, Italy; Department of Public Health, University of Naples Federico II, Naples, Italy; Department of Clinical Physiology, Centre of Postgraduate Medical Education, Warsaw, Poland; National Institute of Cardiology, Warsaw, Poland; Department of Clinical Pharmacology and Toxicology, University Hospital Zurich, University of Zurich, Zurich, Switzerland; Department of Clinical Immunology, Jagiellonian University Medical College, Krakow, Poland

**Keywords:** cardiac fibrosis, heart failure, cardiac fibroblasts, dysferlin, COMP-positive matrifibrocytes, extracellular matrix remodelling

## Abstract

**Background and Aims:** Cardiac fibrosis is a major contributor to heart failure (HF), yet mechanisms limiting pathological fibroblast activation remain incompletely understood. We identified dysferlin (DYSF), a membrane repair protein, as highly induced in HF fibroblasts and investigated its role in regulating profibrotic responses.

**Methods:** Cardiac fibroblasts from patients with end-stage HF and unaffected donor hearts were analysed by liquid chromatography-tandem mass spectrometry and bulk RNA sequencing. Dysferlin expression was validated in independent cohorts. Selected gene/protein expression was validated using single-cell/single-nucleus RNA sequencing and multiplex immunofluorescence of human myocardium from dilated cardiomyopathy (DCM), ischaemic cardiomyopathy (ICM), acute myocardial infarction (AMI), and unaffected hearts. Functional studies were performed in human and mouse cardiac fibroblasts using siRNA-mediated silencing and TGF-β stimulation, and in engineered human 3D cardiac microtissues. Fibrotic remodelling, autophagy, apoptosis, and contractile function were assessed by molecular, histological, biochemical and functional analyses.

**Results:** Dysferlin abundance was markedly increased in HF fibroblasts. Across HF myocardium, DYSF was enriched in activated fibroblasts but largely excluded from COMP-enriched fibrotic regions, consistent with a role in restraining fibroblast state transitions. Although induced by TGF-β, DYSF silencing enhanced extracellular matrix production, increased FOSL2 expression, and promoted differentiation into COMP-positive matrifibrocytes. In engineered human cardiac microtissues, DYSF silencing exacerbated fibrosis, increased apoptosis, and impaired contractility. Mechanistically, dysferlin restrained the TGF-β-FOSL2-autophagy signalling axis, whereas FOSL2 suppressed DYSF expression, defining a reciprocal regulatory circuit. Silencing FOSL2 or MXRA5 increased dysferlin levels, while mRNA-protein discordance implicated S-acylation as a potential regulator of dysferlin protein abundance.

**Conclusions:** Dysferlin is a stress-inducible antifibrotic regulator that limits maladaptive fibroblast differentiation and myocardial fibrosis, thereby representing a potential therapeutic target to attenuate adverse cardiac remodelling in HF.

**Graphical Abstract:** 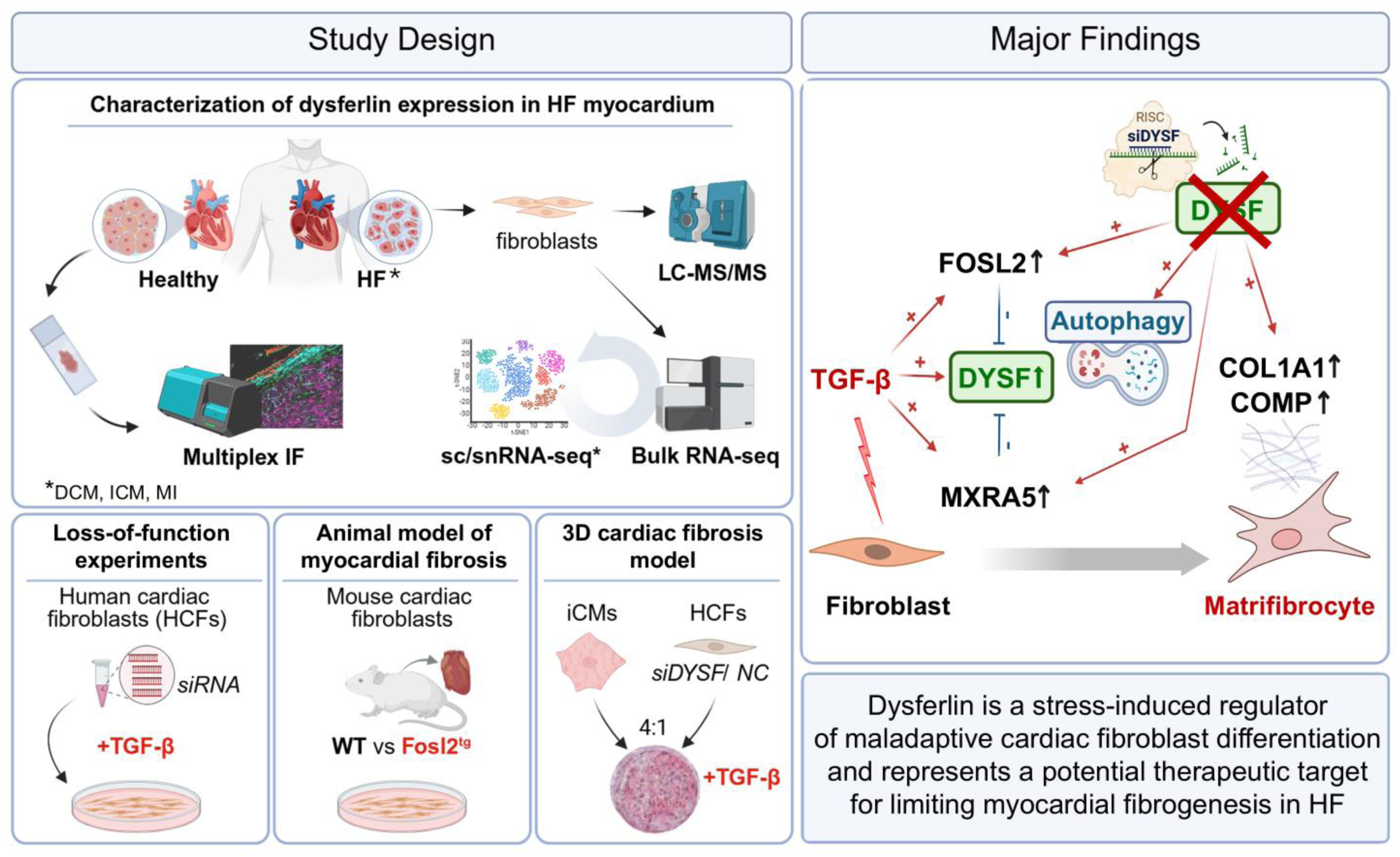

## Introduction

Cardiac fibrosis is characterized by the excessive deposition of extracellular matrix (ECM) proteins in the myocardial interstitium. It represents a convergence point for a broad range of heart diseases and ultimately leads to heart failure (HF) [1]. Pathological stimuli trigger the activation of cardiac fibroblasts, resulting in their proliferation, differentiation, and elevated secretion of ECM molecules. The expanding interstitium increases myocardial stiffness, impairs conduction, and leads to reduced functional performance of the heart muscle. Once initiated, cardiac fibrosis perpetuates its progression, as increasing mechanical stress and a deteriorated tissue environment provide continuous stimuli for fibroblast activation. This persistent activation drives further maladaptive remodeling of the affected myocardium.

The traditional pharmacological management of HF is based on the neurohormonal model of HF pathophysiology, formulated several decades ago [2]. This model attributes HF progression primarily to the activation of the sympathetic nervous system and the renin-angiotensin-aldosterone system (RAAS). As a result, therapeutic interventions - such as beta-blockers or angiotensin-converting enzyme inhibitors - are aimed at suppressing the activation of these neurohormonal pathways. However, recent evidence highlighting the key role of cardiac fibroblast activation challenges the view of cardiac fibrosis as strictly a consequence of neurohormonal activation in HF pathophysiology. Accordingly, novel therapies have shifted the focus from neurohormonal antagonism to more specific cardiac targets [2], [3].

Transforming growth factor β (TGF-β), traditionally considered a master regulator of the fibrotic processes, is known to interplay with other profibrotic mediators and cellular pathways [4]. Fos-like antigen 2 (FOSL2) belongs to the Fos family of transcription factors, which contribute to the formation of the Activator Protein 1 (AP-1) transcription complex. FOSL2 has been implicated in several cellular processes and pathways, including the regulation of TGF-β signalling, and, therefore, is associated with promoting fibrotic changes in multiple organs, including heart [5]–[12]. Classical TGF-β-dependent differentiation of cardiac fibroblast has been linked to increased autophagic activity and can be blocked by autophagy inhibitors *in vitro* [13]. Furthermore, the TGF-β-FOSL2-autophagy axis has been reported to control cardiac fibroblast differentiation, driving the progression of myocardial fibrosis [7].

Rapidly evolving methods of omics technologies have provided a tool for the comprehensive characterization of activated cardiac fibroblasts as fundamental contributors to the fibrotic process, creating a basis for therapeutic targeting of their molecular signature. Recently, several independent research groups have employed single-cell and single-nucleus RNA sequencing (sc/snRNA-seq) to analyse the transcriptomic landscape of fibrotic human myocardium [14]–[19]. These studies revealed substantial heterogeneity among cardiac fibroblast phenotypes, identifying several subsets based on their gene expression profiles. Pathological stimuli further increase this heterogeneity by promoting fibroblast transitions into activated states. Among disease-associated fibroblast subsets, ACTA2-positive myofibroblasts and COMP-positive matrifibrocytes appear to be of particular relevance, with both ACTA2 and COMP upregulated by TGF-β signalling. While myofibroblasts represent a canonical injury-induced activated fibroblast state, matrifibrocytes are a more recently described subset that persists within scarred regions of the fibrotic myocardium.

Regardless of the advantage of uncovering the heterogeneous subsets of cardiac fibroblasts with single-cell precision, transcriptomic datasets alone do not account for post-transcriptional and post-translational regulatory mechanisms, and therefore, may poorly predict actual protein abundances.

In this study, we performed a proteomic analysis of human cardiac fibroblasts and identified the membrane protein dysferlin (DYSF) as a novel molecular candidate upregulated in HF. Dysferlin is best known for its role in sarcolemma resealing and vesicle trafficking in skeletal muscle fibers and is also highly expressed in cardiac muscle, where it contributes to membrane repair in cardiomyocytes exposed to mechanical stress [20].

Despite these established functions, the role of dysferlin in fibroblast activation has remained largely unexplored. Here, we sought to elucidate the contribution of dysferlin to cardiac fibroblast responses under profibrotic conditions and to assess its potential involvement in the regulation of myocardial fibrogenesis in HF.

## Materials and Methods

### Adult cardiac fibroblasts

Cardiac fibroblasts were isolated from the left atria of patients undergoing heart transplantation due to HF associated with ischemic cardiomyopathy and from unaffected hearts of brain-dead donors. Specimens were collected following the procedure approved by the Ethics Committee of the University of Naples Federico II (approval number 79/18) and in compliance with the principles outlined in the Declaration of Helsinki. The characteristics of the patients are summarised in **Table 1**.

**Table 1.** Clinical characteristics of patients. Patients number 1-5 – healthy hearts; patients 6-10 - heart failure.

| Patient | Gender | Age | Ejection fraction | Clinical diagnosis |
| --- | --- | --- | --- | --- |
| 1 | male | 22 |  | No cardiovascular disease |
| 2 | male | 12 |  | No cardiovascular disease |
| 3 | male | 14 |  | No cardiovascular disease |
| 4 | male | 11 |  | No cardiovascular disease |
| 5 | male | 23 |  | No cardiovascular disease |
| 6 | male | 51 | 20% | Ischemic cardiomyopathy, myocardial infarction<br>6 years before, atrial fibrillation, hypertension |
| 7 | male | 46 | 20-25% | Ischemic cardiomyopathy, myocardial infarction<br>5 years before, atrial fibrillation |
| 8 | male | 52 | 15% | Ischemic cardiomyopathy, myocardial infarction<br>9 years before, hypertension |
| 9 | male | 19 | 30% | Dilation of left atrium and left ventricle,<br>mitral regurgitation |
| 10 | male | 44 | 20% | Ischemic cardiomyopathy, myocardial infarction<br>6 years before, diabetes, hypertension |

The cardiac fibroblast isolation procedure has been described previously by Błyszczuk et al.[56]. Tissue fragments were disaggregated by enzymatic digestion with 0.25% trypsin and 0.1% collagenase II (Sigma-Aldrich). Cardiac fibroblasts were isolated using positive selection immunomagnetic cell sorting with anti-fibroblast MicroBeads (MiltenyiBiotec). The sorted cells were cultured *in vitro* in Dulbecco’s Modified Eagle’s Medium (DMEM) supplemented with 10% FBS, penicillin 10,000 U, and streptomycin 10 mg/mL (all from Sigma-Aldrich) at 37 °C in 5% CO_2_. The cells in passages 4-5 were harvested for subsequent transcriptomic and proteomic analysis.

### Proteomic analysis

The cells were resuspended in SDS buffer (4% SDS, 100 mM Tris/HCl pH 8.2, 0.1 M DTT), followed by boiling (95 °C, 5 min) and ultrasonication (1 min). The samples were further processed according to the filter-aided sample preparation (FASP) protocol [57] and purified with StageTips [58]. Liquid chromatography-tandem mass spectrometry (LC-MS/MS) was conducted using Orbitrap Lumos MS instrument (Thermo Fisher Scientific). The protein identification and quantification was performed using MaxQuant (v1.6.2.3) and the *Andromeda* search engine [59], [60]. The data were searched against the human database (fgcz_9606_reviewed_cnl_20190709). A set of functions implemented in the R package SRMService [61] was used to filter proteins quantified with at least two peptides, and the R/Bioconductor package limma [62] was employed to compute moderated t-test [63] for these proteins.

### Bulk RNA sequencing

Total RNA was extracted from the patient’s cardiac fibroblasts using Quick-RNA Microprep Kit (Zymo Research) according to the manufacturer’s instructions. Subsequent RNA sequencing was outsourced to an external service (Genewiz, Germany). Strand-specific RNA sequencing with polyA selection was performed for RNA-seq library preparation. Sequencing was conducted using the Illumina HiSeq platform with paired-end (PE 2×150) configuration. Trimmomatic (v.0.36) was used to trim sequence reads, removing possible adapter sequences and nucleotides with poor quality [64]. Mapping of the trimmed reads was performed to the Homo sapiens GRCh38 reference genome available on ENSEMBL using the STAR aligner (v.2.5.2b) [65]. Unique gene hit counts were calculated using featureCounts [66] from the Subread package (v.1.5.2) [67]. Unique reads falling within exon regions were counted. Gene hit counts were used for downstream differential expression analysis. Gene expression in healthy and HF samples was compared using DESeq2 [68]. The Wald test was applied to compute log2 fold changes and p-values. Genes with an adjusted p-value < 0.05 and absolute log2 fold change > 1 were considered differentially expressed (DEGs). Significant DEGs were clustered by their gene ontology (GO), and GO enrichment was assessed with Fisher’s exact test using the GeneSCF v1.1-p2 tool [69].

### Analysis of publicly available sc/snRNA-seq datasets

The scRNA-seq datasets from human HF published by Wang et al. [17] and deposited in the Gene Expression Omnibus (GEO) under accession numbers GSE109816 and GSE121893 were used to select candidate genes for further analysis. We used processed data and cell clustering described in the original paper to identify DEGs in HF fibroblasts with Seurat package (V.2.3.4) [70]. The obtained DEGs were compared with those identified in our bulk RNA-seq and LC-MS/MS analysis.

The expression of selected candidates was further explored in both healthy (ERP123138 [27]) and HF (SCP1303 [15], GSE183852 [14], EGAS00001006374 [16]) sc/snRNA-seq datasets. The HF samples were restricted to those associated with dilated cardiomyopathy (DCM), while hypertrophic cardiomyopathy samples were excluded from the analysis to ensure greater consistency in the results. The sequencing workflow and data processing methods are described in the original publications. We used processed datasets containing original cell annotations and sample metadata. The datasets were converted to Seurat objects and further analyzed using the Seurat package (v4.4.0).

*DYSF* expression was visualized across myocardial cell types using the original cell annotations and dimensionality reductions provided by the corresponding studies. To compare *DYSF* expression between healthy and DCM samples, sample-level pseudobulk profiles were generated separately for each cell type by summing raw RNA counts across cells or nuclei from the same sample and cell type. Pseudobulk profiles with library sizes below 50,000 counts were excluded. Counts were normalized using edgeR, and log2 counts per million values were calculated. *DYSF* expression was compared between healthy and DCM samples within cardiomyocytes, fibroblasts, and endothelial cells using two-sided Wilcoxon tests.

To investigate matrifibrocyte-associated fibroblast states, the cardiac fibroblast compartment from the Amrute et al. CITE-seq dataset (GSE217494) was reanalysed. Only RNA counts were retained for downstream analysis. A Seurat object was generated using the original sample, cell-type, heart failure aetiology, and other metadata provided by the study. Cells annotated as fibroblasts in the original metadata were subset for further analysis. Fibroblasts were split by sample and normalized using SCTransform, with mitochondrial transcript percentage regressed out. The top 3,000 integration features were selected, and the normalized sample-level datasets were merged. Principal component analysis was performed using 50 principal components, followed by Harmony integration using sample as the integration variable. UMAP dimensionality reduction, nearest-neighbour graph construction, and clustering were performed using the first 30 Harmony-corrected principal components. Clustering was evaluated across resolutions 0.1-0.8, and resolution 0.3 was selected for downstream annotation.

Potential contamination clusters were identified from the initial fibroblast reclustering based on marker gene expression and removed before the final analysis. The remaining fibroblasts were reintegrated with Harmony, reclustered, and annotated into fibroblast subclusters labelled FB1-FB6. Expression of selected markers, including *ACTA2, DYSF, COMP, CILP, CHAD*, and *THBS4*, was visualized on UMAP embeddings and across fibroblast subclusters.

For sample-level comparisons, raw fibroblast counts were aggregated by summing counts across all fibroblasts from each sample. Samples were retained if they had at least 10,000 counts and at least 100 fibroblasts. Counts were normalized using edgeR, and log2 counts per million values were calculated. Differential expression of selected genes was tested using limma linear models with heart failure aetiology as the group variable. Contrasts included ischemic cardiomyopathy, non-ischemic cardiomyopathy, and acute myocardial infarction versus donor controls, as well as pairwise comparisons between disease aetiologies. For the four predefined matrifibrocyte markers *COMP*, *CILP*, *CHAD*, and *THBS4*, Benjamini-Hochberg false discovery rate correction was applied across the marker set within each contrast. DYSF was analysed as an individual candidate gene and is reported using nominal p values.

For the extended analysis of transcriptomic and proteomic results, we used several open-source, web-based tools. Functional enrichment analysis of DEGs was performed using EnrichR [71]. The STRING database with its built-in functional enrichment visualization tool was used to identify the most enriched Gene Ontology (GO) terms for the selected proteins [72]. The interactive platform for Reference of the Heart Failure Transcriptome (ReHeaT) was used to explore the deregulation of the selected genes in HF myocardium across 16 independent studies [73]. Protein palmitoylation datasets were explored using SwissPalm [74].

### *In vitro* cell culture

#### Foetal cardiac fibroblasts

Foetal human cardiac fibroblasts (fHCFs) were purchased from Sigma (Cell Applications) and used for the experiments at passages 10–18 (>99% collagen I-positive, >99% vimentin-positive, and <5% positive for a filamentous form of α-SMA). The cells were cultured in DMEM High Glucose (Sigma-Aldrich) supplemented with 10% FBS (heat-inactivated, Gibco), 1% penicillin/streptomycin (Gibco), and 0.1% 2-mercaptoethanol (50 mM, Gibco).

#### Adult cardiac fibroblasts and endothelial cells

Approximately 1 g of muscle tissue from the left ventricle was transferred into PBS and placed in a 10 cm Petri dish containing 4 mL of complete growth medium (PromoCell Endothelial Cell Growth Medium MV2, Merck), additionally supplemented with 5% human AB serum (Merck), 10 ng/mL HB-EGF (BioLegend), and 100 µg/mL Liberase™ TM (Roche) under sterile conditions. The tissue was cut into pieces smaller than 1 mm using microscissors, and any remaining larger fragments were minced with a scalpel. The tissue was evenly distributed into ten 2 mL Eppendorf tubes, and 1 mL of complete growth medium containing 100 µg/mL Liberase™ TM was added to each tube. The tubes were incubated in a thermomixer at 37 °C and 800 rpm for 1 hour. After digestion, the partially dissociated tissue was transferred into 15 mL tubes, pipetted 10 times using a 10 mL serological pipette, and centrifuged at 350 × g for 5 minutes. The supernatant was removed, 12 mL of PBS was added, and the samples were centrifuged again at 350 × g for 5 minutes. The resulting tissue pellet was transferred into a T75 flask containing 20 mL of complete growth medium and incubated overnight under standard culture conditions. The following day, the medium and tissue were collected into a 50 mL tube and pipetted 15 times with a 10 mL serological pipette to obtain a single-cell suspension containing microvascular fragments. The suspension was centrifuged at 300 × g for 5 minutes, and the cells were seeded into culture flasks pre-coated with 0.2% porcine gelatin (Merck).

Cells were cultured for up to 6 days or until 70% confluence was reached, then harvested using 0.25% trypsin–EDTA solution (Merck). The cells were centrifuged, resuspended in PBS, and labeled for 10 minutes with anti-CD31-biotin (clone WM-59, Thermo Fisher Scientific). Cells were magnetically sorted using the CELLection™ Biotin Binder Kit (Thermo Fisher Scientific) according to the manufacturer’s instructions. CD31-negative adult human cardiac fibroblasts (adHCFs) were cryopreserved in growth medium containing 10% DMSO (Merck).

CD31-positive endothelial cells (ECs) were cultured to full confluence with medium changes three times per week and passaged at a 1:3 ratio. Second-passage ECs were cryopreserved in growth medium containing 10% DMSO (Merck). After thawing, >95% of the cells were CD31- and von Willebrand factor-positive. Isolations were performed following approval from the Bioethical Committee (approval no. IK.N.PIA.002.4.2128/25).

For the experiments, adHCFs were cultured in DMEM/F12 (Sigma-Aldrich) supplemented with 10% FBS, 1% penicillin/streptomycin, and 0.1% 2-mercaptoethanol (50 mM, all from Gibco). Fibroblast activation was induced by stimulation with TGF-β (10 ng/mL, PeproTech) for 24–72 hours. TGF-β receptor I (TGF-βRI) was inhibited using the selective inhibitor SD208 (100 nM, Tocris). ECs were seeded on plates coated with 0.2% gelatin and cultured in Endothelial Cell Growth Medium MV2 (Sigma-Aldrich) supplemented with 5% human serum (Sigma-Aldrich) and 10 ng/mL HB-EGF (BioLegend).

#### iPSC-derived cardiomyocytes (iCMs)

Differentiated, contracting iCMs for 2D monoculture experiments were obtained from the iPSC Core Facility at the Institute for Regenerative Medicine, University of Zurich. These cells were cultured in maintenance medium composed of DMEM High Glucose (Sigma-Aldrich), 2% FBS (Gibco), 50 µM phenylephrine hydrochloride (Sigma-Aldrich), 0.3 µM L-ascorbic acid (Sigma-Aldrich), 1% penicillin/streptomycin (Gibco), and 0.1% 2-mercaptoethanol (50 mM, Gibco). The iCMs were passaged using the Multi Tissue Dissociation Kit 3 (Miltenyi Biotec) in accordance with the manufacturer’s instructions. Dissociated iCMs were seeded on plates coated with Synthemax II-SC Substrate (Corning).

For palmitoylation inhibition experiments, ECs were seeded in 24-well plates at a density of 30,000 cells/well and iCMs in 12-well plates at 200,000 cells/well. Cells were treated with 25–50 µM 2-bromopalmitate (Sigma-Aldrich) for 24 hours. Whole-cell lysates were collected for subsequent Western blot analysis.

### Gene silencing

The cells were transfected using Lipofectamine 2000 (Thermo Fisher Scientific) according to the manufacturer’s instructions. Predesigned siRNA directed against human *DYSF* (GS8291), *MXRA5* (GS25878), and *FOSL2* (GS2355) were purchased in the form of FlexiTube GeneSolution kits (Qiagen). Non-targeting siRNA was used as a negative control (Qiagen). The cells were transfected under antibiotic-free conditions with a 50 nM siRNA solution. After 6 hours, the culture medium containing the transfection mix was replaced with fresh culture medium.

### Cell viability and apoptosis assays

For the resazurin-based viability assay, fHCFs, adHCFs and ECs were seeded in 24-well plates at a density of 30,000 cells/well, and iCMs were seeded in 12-well plates at 200,000 cells/well. The assay was performed 48 h after transfection. The cells were incubated with PrestoBlue HS reagent (Invitrogen) added to the culture medium at a 1:10 ratio for 3 hours in a cell incubator (37°C, 5% CO₂). Next, the supernatants were transferred to a 96-well plate, and the conversion of resazurin to fluorescent resorufin was measured using the Synergy HT microplate reader (BioTek). As this is a live assay, the remaining cells were used for further RNA extraction and RT-qPCR.

fHCFs seeded onto a 96-well plate at 4,000 cells/well were used for measuring ATP levels and apoptotic activity. These parameters were quantified using CellTiter-Glo and Caspase-Glo 3/7 assays (Promega), respectively. The procedures were carried out in accordance with the manufacturer’s instructions. Following incubation with the assay reagents, the plates were directly used for luminescent readout with the Synergy HT (BioTek).

### Contraction assay

To examine the contractile properties of fHCFs, we used the Contraction Assay Kit (Cell Biolabs) following the manufacturer’s protocol. The cells were preliminarily transfected either with the appropriate siRNA against a selected candidate or with non-targeting control siRNA, as described above. Next, the cells were trypsinized and resuspended in medium at a concentration of 4 million cells/mL. The gels were prepared by mixing 50 µL of cell suspension and 200 µL of collagen mixture per gel, which resulted in 200,000 cells/gel in a well of a 48-well plate. Complete culture medium containing 20 ng/mL of TGF-β was applied on top of the solidified gels.

After 48 hours of incubation, the gel matrices were released from the walls of the culture plate (time point 0), and images were taken at this and subsequent time points up to 72 hours. The pictures were taken using the Fusion FX (Vilber) instrument. The areas of the gels were measured using ImageJ and normalized to those at time point 0.

### RT-qPCR

Spin-column purification of total RNA was performed using the Quick-RNA MicroPrep Kit (Zymo Research). The culture medium was removed from the plate, and the cells were lysed with the provided RNA lysis buffer (Zymo Research). Next, the lysates were mixed with an equal volume of ethanol (≥99.8%, Honeywell) and transferred onto Zymo-Spin IC Columns. Further centrifugation and washing steps were performed as described in the manufacturer’s protocol, including on-column DNase I treatment to remove contaminant DNA. In the final step, RNA was eluted from the column matrix with DNase/RNase-free water (Zymo Research). The concentration and purity of the extracted RNA were quantified using the NanoDrop 2000 (Thermo Fisher Scientific).

The reverse transcription of RNA (150 ng/sample) was performed using MultiScribe Reverse Transcriptase, random hexamer primers, and RNase inhibitor (all Thermo Fisher Scientific). The complete master mix was prepared according to the manufacturer’s instructions, and the reaction was run in the Bio-Rad T100 Thermal Cycler.

RT-qPCR was then performed using the GoTaq qPCR Master Mix (Promega) on an Agilent Technologies Stratagene Mx3005P instrument. The specific human and mouse primers used for the reaction are listed in **Tables 2 and 3**, respectively. The obtained Ct values were normalized using an endogenous control (housekeeping genes: RPLP0, GAPDH), and relative quantification of gene expression was performed using the 2^−ΔΔCt method [75]. The data are presented as a fold change (FC) relative to the mean of the control group.

**Table 2.**
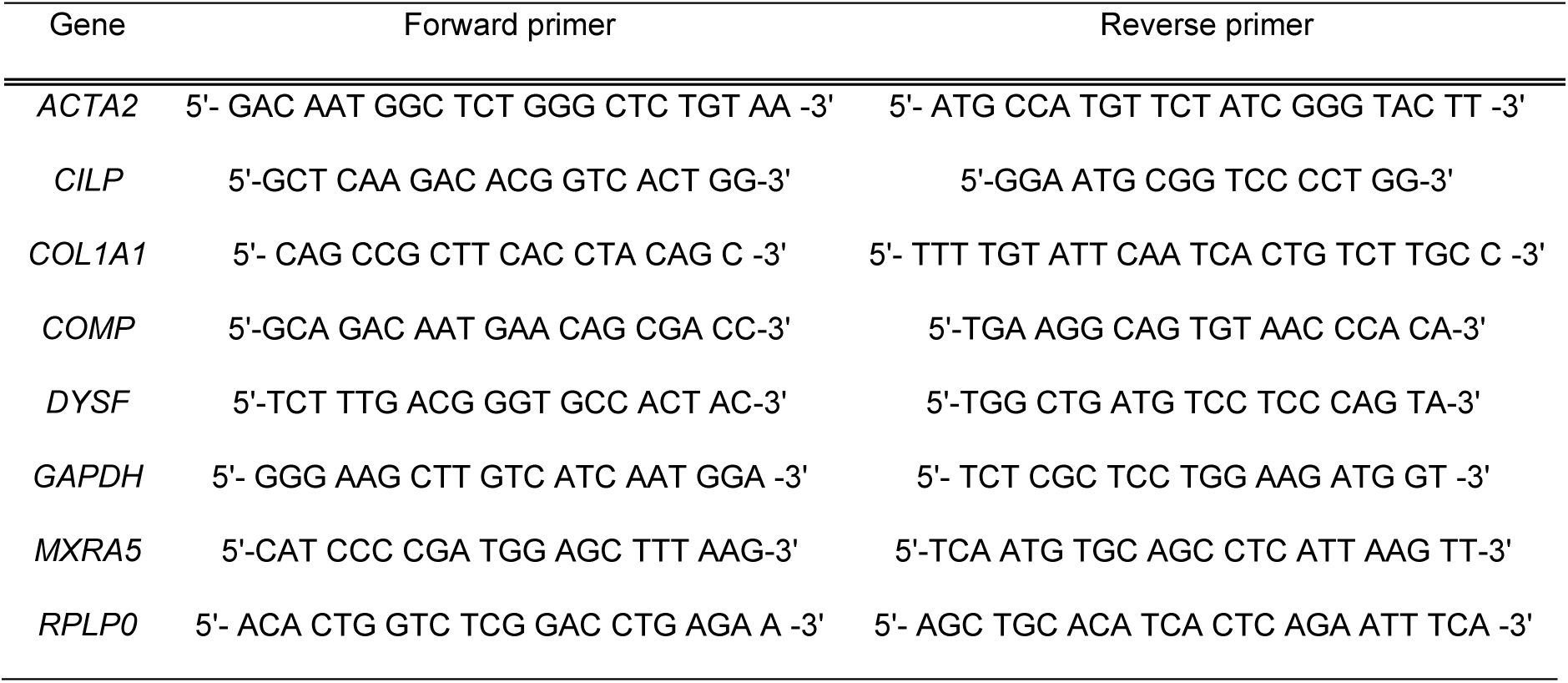
Human primer sequences used for RT-qPC.

**Table 3.**
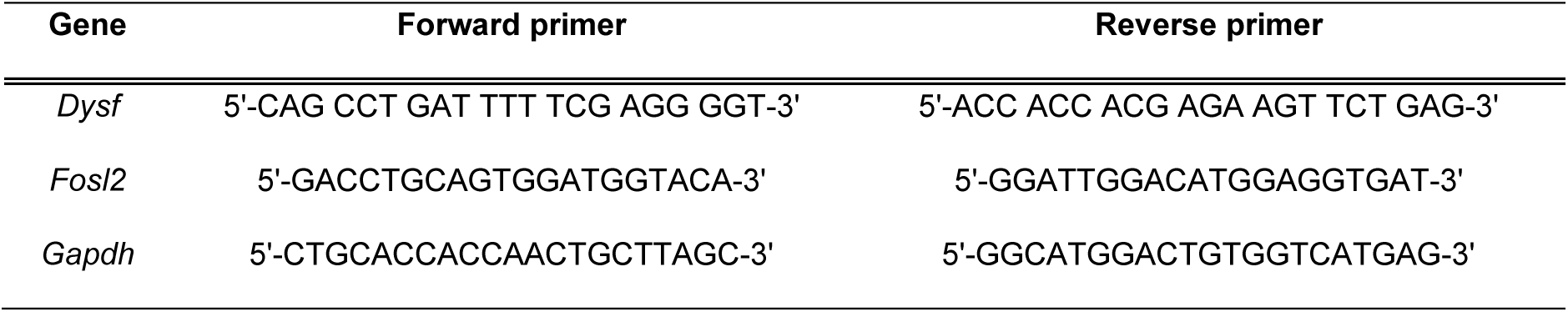
Mouse primer sequences used for RT-qPCR.

### Western blotting

Cells were lysed in RIPA buffer (Sigma-Aldrich) supplemented with phosphatase inhibitors (PhosStop, Roche) and protease inhibitors (cOmplete ULTRA, Roche). Protein concentration in cell lysates was quantified using the Pierce BCA Protein Assay Kit (Thermo Scientific). For SDS-PAGE, cell lysates containing equal amounts of protein were mixed with 10x Reducing agent and 4x LDS sample buffer (both from Invitrogen), followed by boiling at 95°C for 5 minutes. Protein samples prepared this way were loaded onto the SDS-PAGE gel and separated by electrophoresis, followed by overnight wet transfer (30 V, 4°C) onto a nitrocellulose blotting membrane (pore size 0.45 µm, Amersham Protran).

The membranes were reversibly stained with Ponceau S (Sigma-Aldrich) to confirm transfer efficiency by visualizing protein bands. Non-specific binding sites were blocked with 5% skim milk (BD Life Sciences) in TBST solution, and membranes were incubated with primary antibody solution overnight at 4°C. Subsequent incubation with secondary antibodies was conducted for 1 hour at RT. The primary and secondary antibodies used for immunoblotting are listed in **Table 4**.

**Table 4.**
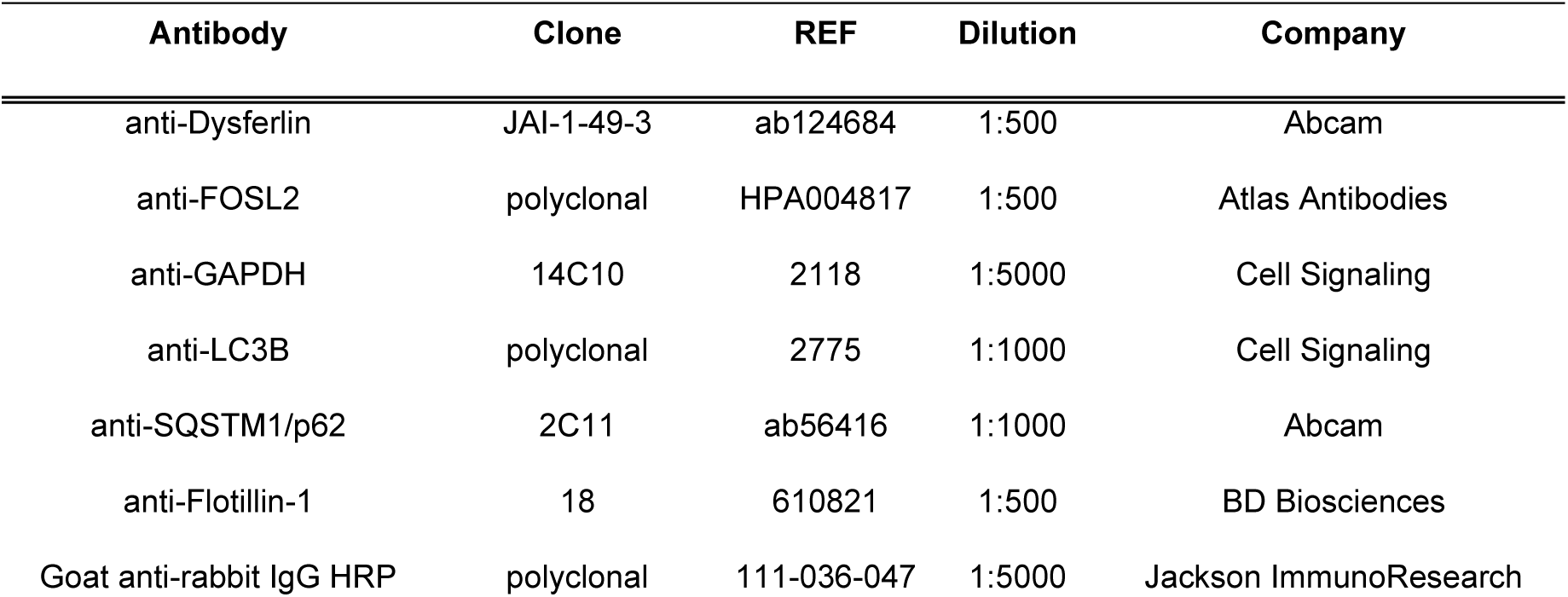
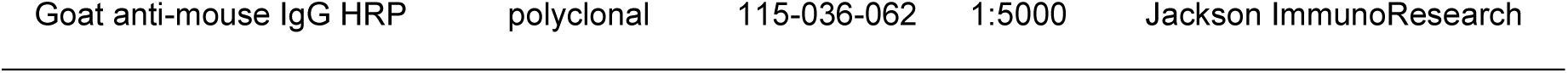
List of antibodies used for Western blotting.

The signal was developed using chemiluminescent substrate (SuperSignal West Pico PLUS, Thermo Scientific) and visualized with the Fusion FX (Vilber) instrument. The acquired signal was quantified by densitometry analysis using ImageJ software. Normalization was performed using a housekeeping protein (GAPDH) as an internal loading control. The results are presented as a fold change (FC) of normalized values relative to the mean of the control group.

### ELISA

Quantification of secreted human pro-collagen-Iα1 was performed using a DuoSet ELISA kit (DY6220-05, R&D Systems) according to the manufacturer’s protocol. The 96-well plates were incubated with capture antibodies overnight at RT, followed by blocking with 1% BSA in PBS. The washing steps were performed using 0.05% Tween 20 in PBS. The supernatants, unconditioned culture medium, and standards were then incubated for 2 hours at RT. After subsequent washing, plates were treated with biotinylated detection antibodies for 2 hours. The consecutive incubation steps with streptavidin-HRP and TMB substrate (BD Biosciences) were carried out for 20 minutes each, protecting the plates from direct light. The reaction was stopped using 2N H₂SO₄ solution. The optical density of each well was measured immediately using the Synergy HT (BioTek) microplate reader set to 450 nm, with wavelength correction at 570 nm. Duplicate measurements were taken for each sample, and concentrations were determined based on the respective standard curves. Pro-collagen-Iα1 concentration in the samples was determined by interpolating the standard curve. The results are presented as a concentration fold change (FC) to the mean of the control group.

### Cardiac microtissue fabrication and analysis

Cardiac microtissues were fabricated from human iCMs and fHCFs mixed in a 4:1 ratio following the procedure described by Blyszczuk et al. [56]. Frozen iCMs were purchased from Cellular Dynamics International (ref. C1058) and used for microtissue assembly directly after thawing. The fHCFs had been transfected with either DYSF or NC siRNA 24 hours in advance (**Scheme 1**). The iCM:fHCF or fHCF-only suspension in maintenance medium was seeded on Akura 96 spheroid microplates (InSphero) at a concentration of 5000 cells per well and kept in a cell culture incubator (37°C, 5% CO₂) for 72 hours to allow for self-aggregation of the contained cells. Subsequently, the assembled iCM:fHCF microtissues and fHCF spheroids were treated with 20 ng/mL TGF-β to simulate fibrotic conditions or maintained as a control (day 0). On day 6 of TGF-β stimulation, the microtissues were collected for downstream analyses.

Total RNA was isolated from 10–12 pooled microtissues using the Quick-RNA MicroPrep Kit (Zymo Research), and RT-qPCR was performed as described above. Apoptotic activity was evaluated using the Caspase-Glo 3/7 assay (Promega), following the manufacturer’s protocol.

Microtissue contraction analysis was performed 72 hours after microtissue assembly (day 0) and repeated on day 6 of TGF-β stimulation. Microtissues were visualized using the AxioObserver Z1 microscope (Zeiss) equipped with a humidified chamber at 37°C and 5% CO₂. Videos of contracting microtissues were recorded using ZEN software and processed with Fiji (ImageJ) in combination with a custom-made macro. Quantitative analysis of microtissue contraction was subsequently conducted with the MUSCLEMOTION open-source tool [76].

**Scheme 1.**
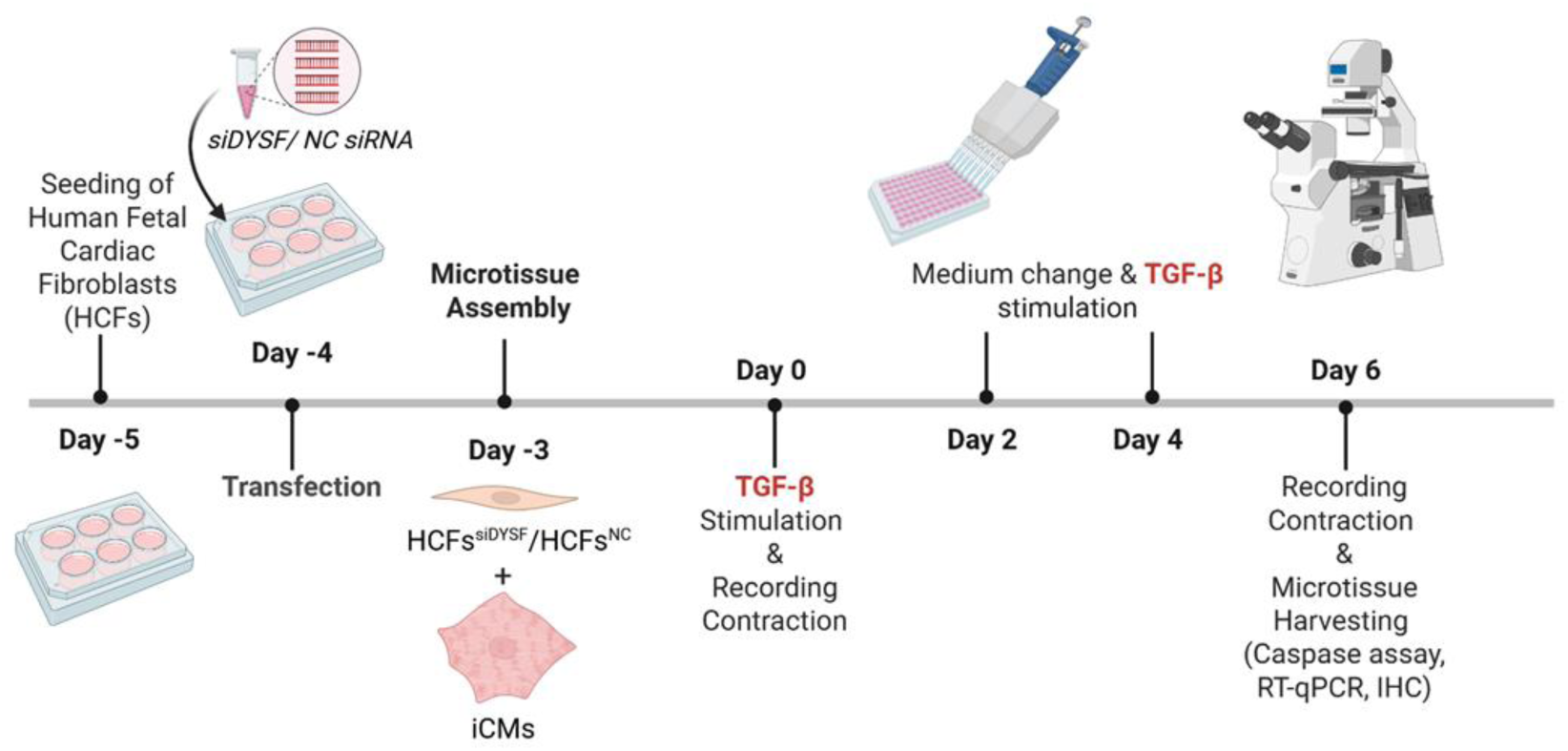
Timeline of cardiac microtissue fabrication and analysis.

### Animal model

Fosl2-overexpressing (Fosl2^tg^) mice were obtained from Sanofi Pharmaceuticals and backcrossed with C57Bl/6 mice (Charles River) for at least 10 generations [5], [7], [8]. Housing and breeding were carried out under pathogen-free conditions at the Laboratory Animal Services Center, University of Zurich. The genotype was confirmed by EGFP expression in blood samples. Animal experiments were performed in accordance with Swiss federal law and approved by the Cantonal Veterinary Office Zurich (approval ZH007/2019).

Fosl2^tg^ mice and wild-type (WT) littermates aged 16 to 20 weeks were used for heart harvesting and subsequent cardiac fibroblast isolation. Following perfusion with ice-cold phosphate-buffered saline (PBS) through the left ventricle, the hearts were collected on ice until further processing. Mechanical and enzymatic dissociation was performed in a 0.025 mg/mL Liberase TM (Collagenase I and II, Roche) solution for 45 min at 37 °C. The resulting cell suspension was passed through a 70 µm cell strainer, followed by centrifugation at 50 × g for 2 min to remove cardiomyocytes. The supernatant was then passed through a 40 µm cell strainer and centrifuged at 300 × g for 4 min. The obtained pellet was resuspended in DMEM high glucose (Sigma-Aldrich) containing 10% FBS, 50 U/mL penicillin, 50 µg/mL streptomycin, and 50 nM β-mercaptoethanol (all from Gibco) and seeded in T-25 culture flasks to allow for fibroblast adhesion. A cultured homogeneous population of fibroblasts was used for WB and RT-qPCR analysis.

**Scheme 2.**
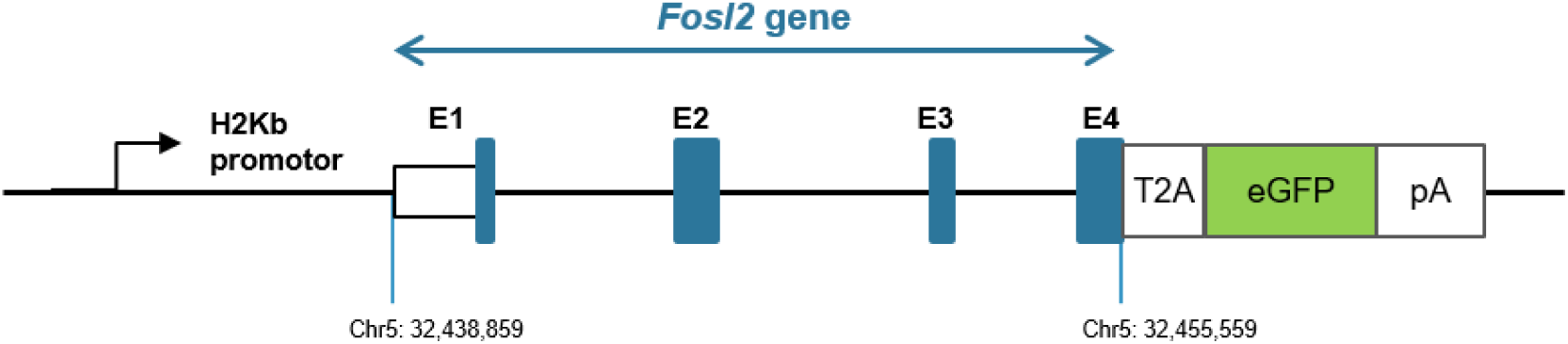
Structure of the genetic construct used for the generation of the Fosl2^tg^ mouse model. A vector comprising exons 1–4 (E1–E4) of the mouse Fosl2 gene was inserted under the control of the H2Kb promoter. The eGFP coding sequence was inserted as a tracer in frame with the Fosl2 gene. A Thosea asigna virus (T2A) self-cleaving peptide was inserted between the Fosl2 and eGFP sequences to induce cleavage of the amino acid chain during translation and prevent Fosl2-eGFP protein fusion. Adapted from Renoux et al., Cell Reports, 2020.

### Immunohistochemistry

Myocardial tissue samples and cardiac microtissues were fixed in 4% paraformaldehyde (PFA, ChemCruz) in PBS, followed by washing in distilled water. Cardiac microtissues were pooled (20-24 per sample) and embedded in 1% agarose prior to further processing. Samples underwent gradual dehydration by incubation three times in 80% ethanol for 1 hour, three times in 96% ethanol for 1 hour, and three times in 100% ethanol for 1 hour, followed by ethanol replacement through three 1-hour incubations in xylene. Subsequently, samples were incubated in paraffin at 56 °C for 3 hours. Sections of 2.5–3 µm thickness were cut, mounted on the Superfrost Plus microscope slides (Thermo Scientific), and dried overnight at 58 °C. Deparaffinization was performed twice in xylene for 3 minutes, followed by rehydration through a descending alcohol series (100%, 100%, 96%, 80%; 3 minutes each). After rehydration and washing in distilled water, antigen retrieval was carried out using Tris-EDTA buffer (pH 9) or citrate buffer (pH 6). Immunodetection was performed on blocked sections (10% goat serum, Vector Laboratories) using the following primary antibodies: anti-dysferlin (1:4000, NBP3-43895, clone CL10888), anti-vimentin (1:10,000, ab92547, clone EPR3776), and anti-COMP (myocardial sections: 1:500, Invitrogen, PA5-95547, polyclonal; cardiac microtissues: 1:8000, BioRad, MCA1455G, clone HC484D1), followed by detection using the Bond Polymer HRP Refine Detection Kit (Leica). Nuclei were counterstained with hematoxylin (J.T.Baker). Stained sections were analyzed using an Olympus BX53 microscope equipped with a camera and cellSens Standard imaging software (Olympus).

### Multiplex sequential immunofluorescence

Formalin-fixed, paraffin-embedded (FFPE) tissue sections (5 µm) were baked in a dry oven for 1 h at 60 °C. Sections were deparaffinized in xylene (5 min, 2 times), followed by incubation in 100% ethanol (2 min, 2 times), and dried for 5 min at 60 °C. Antigen retrieval was performed using the PT Module (Epredia) with HIER Buffer H (pH 9) at 99°C for 1 h. Slides were subsequently washed with Multistaining Buffer (MSB, Lunaphore) and incubated in 5% horse serum (Vector Laboratories) diluted in MSB for 30 min. Next, the slide fitted with a COMET Chip, and the prediluted antibodies (**Table 5**) were loaded into the COMET instrument (Lunaphore) according to the manufacturer’s instructions. Automated sequential staining, imaging, and elution cycles were performed as previously described [77]. The resulting multiplex image stack was exported as an OME-TIFF file and analyzed using HORIZON Viewer software (Lunaphore).

**Table 5.**
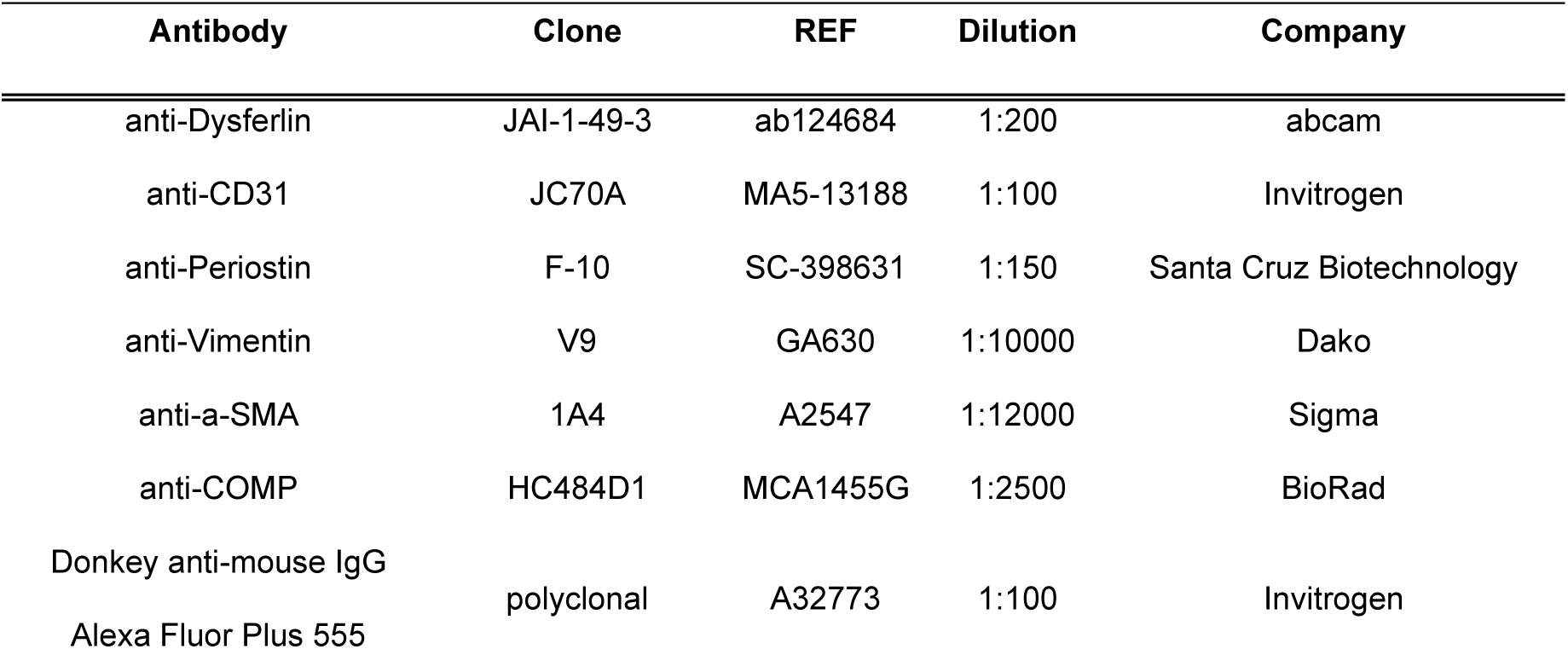
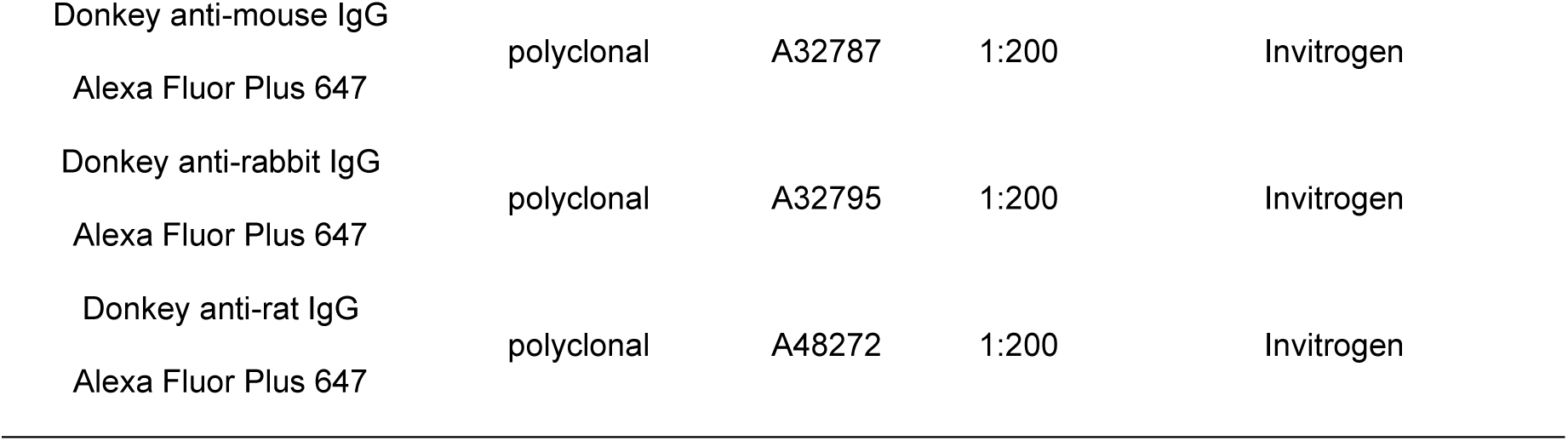
List of antibodies used for sequential Immunofluorescence.

### Statistics

Statistical analyses of experimental data were performed using GraphPad Prism 10 software. The Robust Regression and Outlier Removal (ROUT) method, with a false discovery rate below 1%, was used to identify outliers. The normality of data distribution was examined using the Shapiro-Wilk test. For two-group comparisons, normally distributed data were analysed using an unpaired t-test, while non-normally distributed data were analysed using the unpaired non-parametric Mann-Whitney U test. For multiple group comparisons with one or two variables, analysis was performed using one-way or two-way ANOVA, respectively. Tukey’s or Sidak’s post hoc tests were performed to adjust for multiple comparisons. The analyses were performed with the GraphPad Prism 10 software. The statistical tests are specified in the figure legends.

## RESULTS

### 1. Dysferlin is upregulated in HF-derived cardiac fibroblasts

Human cardiac fibroblasts (HCFs) isolated from the left atria of patients with end-stage HF (n=5) and from unaffected donor hearts (n=5) were subjected to liquid chromatography-tandem mass spectrometry (LC–MS/MS) to identify alterations in their proteomic profiles. This analysis revealed 14 differentially abundant proteins in HF-derived fibroblasts compared with healthy controls (**Figure 1a-c**). Functional enrichment analysis using Enrichr indicated that deregulated proteins were predominantly associated with ECM and membranous vesicles (**Supplementary Figure 1a**), involved in processes related to ECM remodelling (**Supplementary Figure 1b**), and linked to several pathological conditions, such as myocardial ischemia (**Supplementary Figure 1c**).

**Figure 1.**
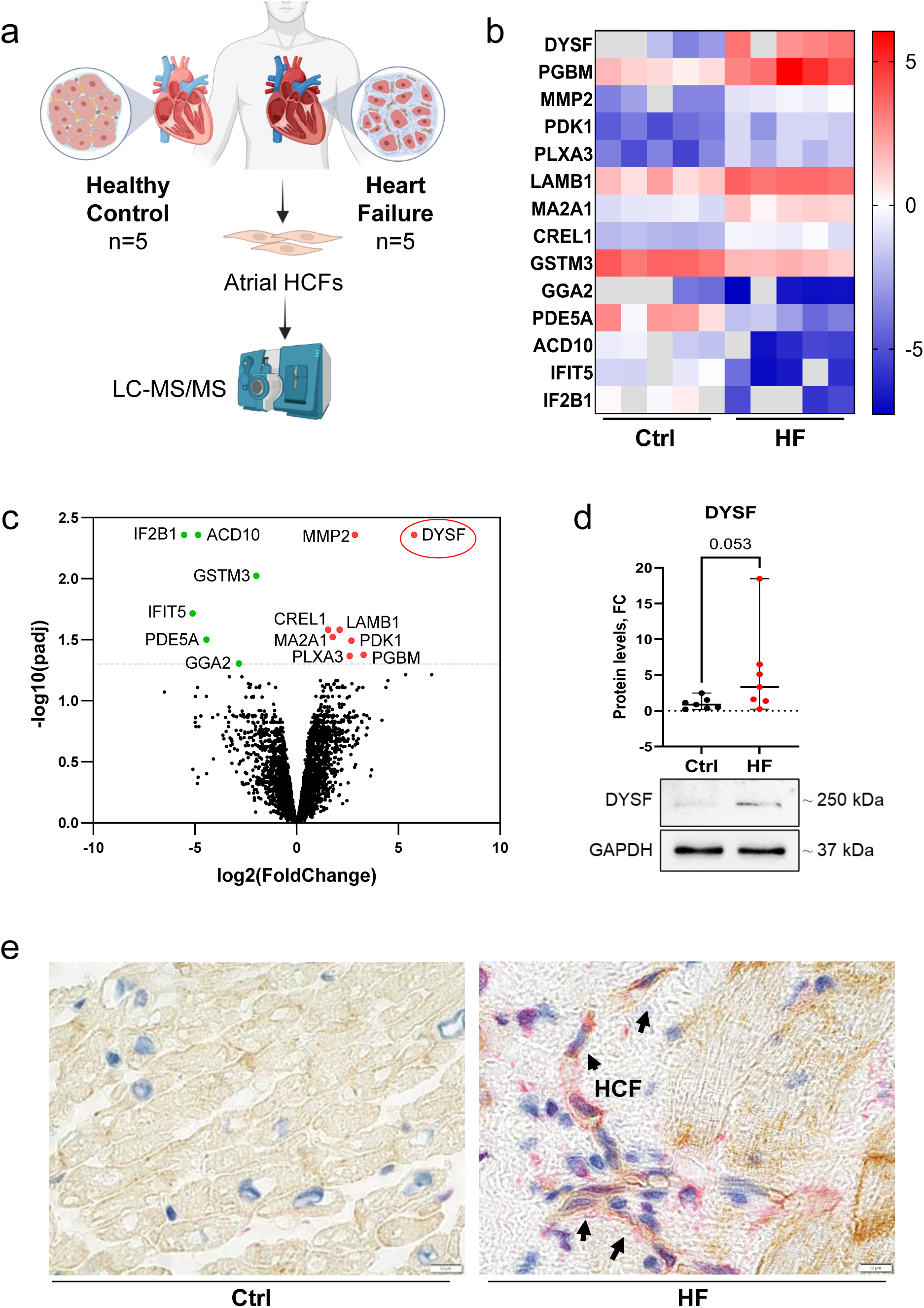
Dysferlin is upregulated in HF-derived cardiac fibroblasts. **(a)** Schematic representation of the proteomic analysis of human cardiac fibroblasts (HCFs) isolated from the left atria of donors and HF patients. Protein quantification was performed using liquid chromatography-tandem mass spectrometry (LC-MS/MS). The image was created with BioRender.com. **(b)** Heatmap and **(c)** volcano plot illustrating differentially expressed proteins (absolute log₂FC > 1, adj. p < 0.05) in the HF group compared with healthy controls. In the volcano plot, red and green dots indicate proteins significantly upregulated and downregulated in HF, respectively. The most upregulated protein, dysferlin, is highlighted in red in the volcano plot. **(d)** Representative immunoblots and corresponding quantification of dysferlin levels in left ventricular fibroblasts from an independent cohort of donors (Ctrl) and HF patients (n=7; Mann-Whitney U test; median ± 95% CI). **(e)** Representative images of IHC staining for dysferlin and vimentin in left ventricular tissue sections from donors (Ctrl) and HF patients. Arrows indicate HCFs. Scale bars: 10 µm.

Dysferlin emerged as one of the most upregulated proteins in HF-HCFs (DYSF, log2FC=5.78, adjusted p < 0.004) and was selected for further validation as a potential candidate associated with myocardial fibrogenesis in HF. Functional exploration of dysferlin using the STRING database and enrichment analysis revealed its involvement in plasma membrane-related processes and components, including plasma membrane repair, T-tubule organization, and vesicle fusion (**Supplementary Figure 2a-b**). Furthermore, the top enriched biological processes reflected the established role of dysferlin in skeletal muscle development and function (**Supplementary Figure 2a**). Enrichment of molecular functions related to calcium-dependent phospholipid and protein binding, as well as calcium ion binding, highlights dysferlin’s role as a Ca²⁺ sensor and effector, consistent with its function in Ca²⁺-triggered vesicle fusion during membrane repair (**Supplementary Figure 2c**).

Dysferlin upregulation in HF was subsequently validated using HCFs isolated from the left ventricles of an independent cohort of DCM patients (n=7) and donor hearts (n=7). The immunoblotting results demonstrate upreguated dysferlin protein levels in HF-derived ventricular HCFs (log2FC=1.94, p=0.053) (**Figure 1d**). Immunohistochemical (IHC) staining of left ventricular myocardial sections revealed an increased number of fibroblasts co-expressing vimentin (VIM) and dysferlin, as well as more pronounced dysferlin expression in HF (**Figure 1e, Supplementary Figure 3**). Dysferlin expression was most prominent at the cardiomyocyte plasma membrane, particularly in regions corresponding to the intercalated discs (ICDs). Notably, ICDs in HF exhibited a disrupted and irregular architecture (**Supplementary Figure 3**), consistent with previous reports describing structural remodelling of ICDs in cardiac disease [21], [22]. Together, these observations suggest that dysferlin expression is induced under pathological conditions associated with HF.

### 2. Dysferlin is a TGF-β-induced negative regulator of profibrotic activation

Following the proteomic identification of dysferlin as an upregulated protein in HF fibroblasts, we performed a series of *in vitro* experiments to unravel its specific role and expression pattern in cardiac fibroblasts under profibrotic conditions. Using foetal human cardiac fibroblast (fHCF) *in vitro* culture, we demonstrated that TGF-β stimulation upregulated dysferlin at both mRNA and protein levels (**Figure 2a-b**). This effect was abolished by pharmacological inhibition of TGF-β receptor I (TGF-βR1) using SD208 (**Figure 2b**).

**Figure 2.**
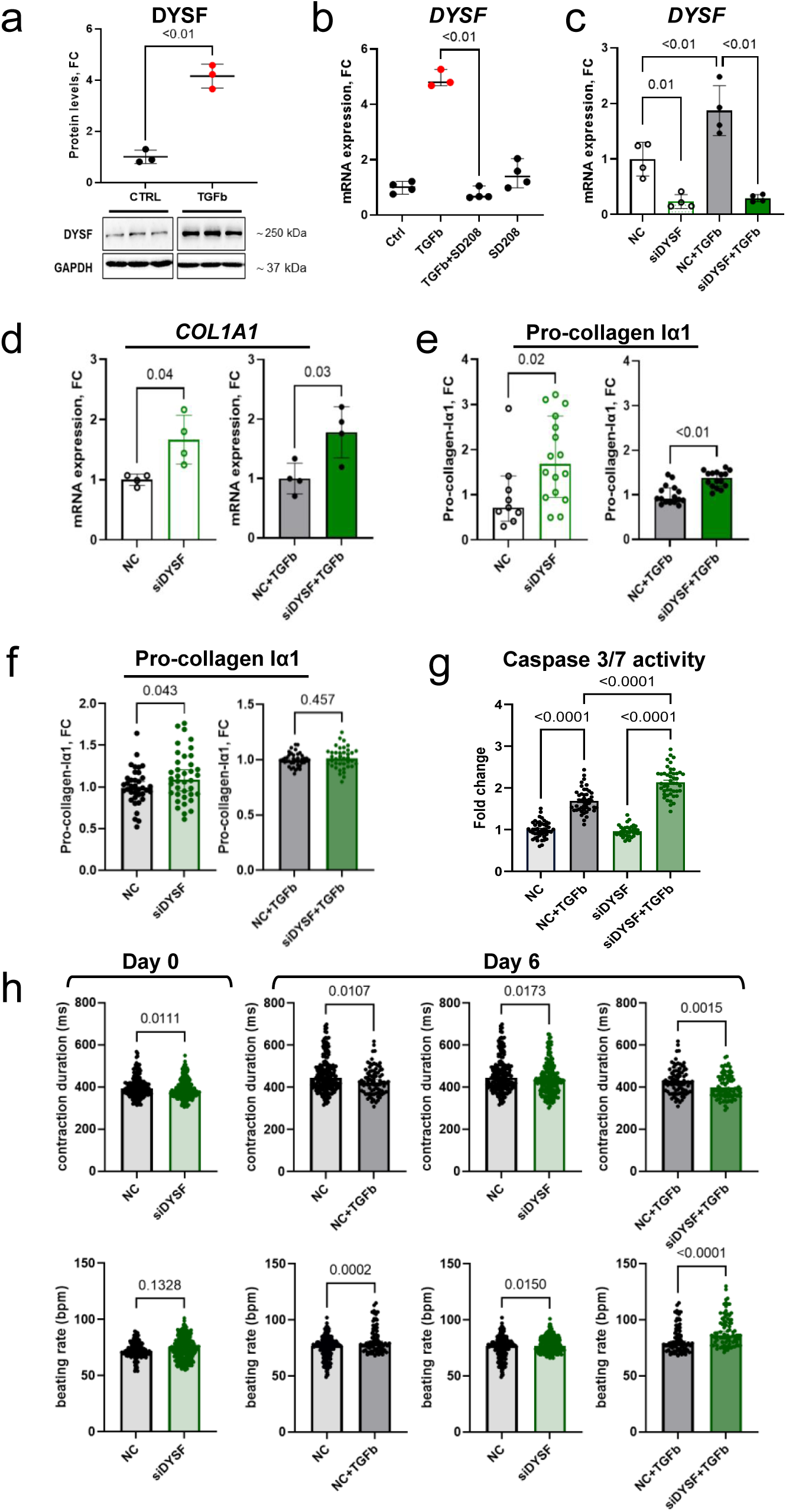
Dysferlin is a TGF-β-induced negative regulator of profibrotic activation. **(a)** Quantification of dysferlin protein levels in cultured fHCFs following 72 h of TGF-β stimulation (10 ng/mL) and corresponding immunoblots (n=3; unpaired t-test; mean ± SD). **(b)** Quantification of *DYSF* gene expression following 48 h of TGF-β stimulation (10 ng/mL) in the presence or absence of the selective TGF-βRI (ALK5) inhibitor SD208 (1 µM), assessed by RT-qPCR (n=3-4; Kruskal-Wallis test, Dunn’s multiple comparisons test; median ± 95% CI). **(c)** *DYSF* gene expression in fHCFs transfected with either *DYSF*-targeting siRNA (siDYSF) or negative control (NC) siRNA, 48 h after TGF-β stimulation (10 ng/mL) (n=4; one-way ANOVA, Tukey’s multiple comparisons test; mean ± SD). **(d)** *COL1A1* gene expression in transfected fHCFs (n=4; unpaired t-test; mean ± SD). **(e)** Levels of pro-collagen Iα1 secreted by 3D fibroblast spheroids composed of transfected fHCFs after 48 h of TGF-β stimulation, measured by ELISA in culture supernatants (n=9-16; Mann-Whitney U test; median ± 95% CI). **(f)** Secreted pro-collagen Iα1 from 3D cardiac microtissues stimulated with TGF-β (20 ng/mL, 6 days) or cultured under control conditions (n=37-40). **(g)** Apoptotic activity in cardiac microtissues after 6 days of TGF-β treatment (20 ng/mL), measured using the Caspase-Glo 3/7 assay. Each value represents one microtissue (n=30-32; one-way ANOVA, Tukey’s multiple comparisons test; mean ± SD). **(h)** *DYSF* knockdown combined with TGF-β stimulation (20 ng/mL for 6 days) alters the contractile properties of cardiac microtissues. Contraction parameters, including contraction duration (ms) and beating rate (bpm), are shown for each recorded microtissue (n=84-207). Statistical analysis for panels **(f)** and **(h)**: two-tailed parametric t-test (mean ± SEM) for normally distributed data and Mann-Whitney U test (median ± 95% CI) for non-normally distributed data. NC: negative control; siDYSF: *DYSF* silencing.

Next, we performed the loss-of-function experiments to investigate the effects of siRNA-mediated *DYSF* knockdown in fHCFs. *DYSF* silencing resulted in upregulated *COL1A1* expression in both untreated and TGF-β-stimulated fHCFs cultured in a monolayer (**Figure 2c-d**). This finding was further supported by results from 3D-cultured fHCFs, in which *DYSF* silencing increased secreted levels of pro-collagen- IA1 under both control and TGF-β-stimulated conditions, indicating that dysferlin loss enhances the profibrotic activation of cardiac fibroblasts (**Figure 2e**).

Next, we used the 3D cardiac microtissue model to validate the role of dysferlin on a higher level of biological relevance in the context of myocardial tissue function. The microtissues were composed of human induced pluripotent stem cell-derived cardiomyocytes (iCMs) and fHCFs. Prior to assembly, fHCFs were transfected with either *DYSF*-targeting (siDYSF) or NC siRNA. The spontaneously contracting microtissues were then maintained under control conditions or treated with TGF-β for 6 days to induce a fibrotic phenotype, confirmed by Masson’s trichrome staining (**Supplementary Figure 4**). Under control conditions, siDYSF microtissues increased secreted levels of pro-collagen Iα1 compared with the corresponding NC microtissues (**Figure 2f**). Under profibrotic conditions, *DYSF* silencing in fHCFs exacerbated TGF-β-induced apoptotic activity in cardiac microtissues, as shown by caspase-3/7 assay (**Figure 2g**).

Furthermore, we assessed the contractile properties of cardiac microtissues. Prior to TGF-β stimulation, *siDYSF* microtissues (day 0) exhibited decreased contraction duration and a trend toward increased beating rate (**Figure 2h**). By day 6, microtissues comprising *DYSF*-silenced fHCFs displayed a higher beating rate and shorter contraction duration compared with NC microtissues under both control and TGF-β–stimulated conditions, with more pronounced effects upon profibrotic stimulation (**Figure 2h**).

Overall, *DYSF* knockdown in fHCFs enhanced the profibrotic phenotype in both fibroblast monoculture and cardiac microtissues, suggesting that dysferlin may act as a negative regulator of profibrotic crosstalk.

### 3. *DYSF* silencing promotes a matrifibrocyte-like phenotype in HCFs

To further investigate the functional consequences of *DYSF* silencing in cardiac fibroblasts, we performed a collagen gel contraction assay using transfected fHCFs embedded in a three-dimensional collagen matrix and cultured under profibrotic conditions. Gels containing *DYSF*-silenced fHCFs exhibited reduced contractility compared with negative control (NC) cells (**Figure 3a**).

**Figure 3.**
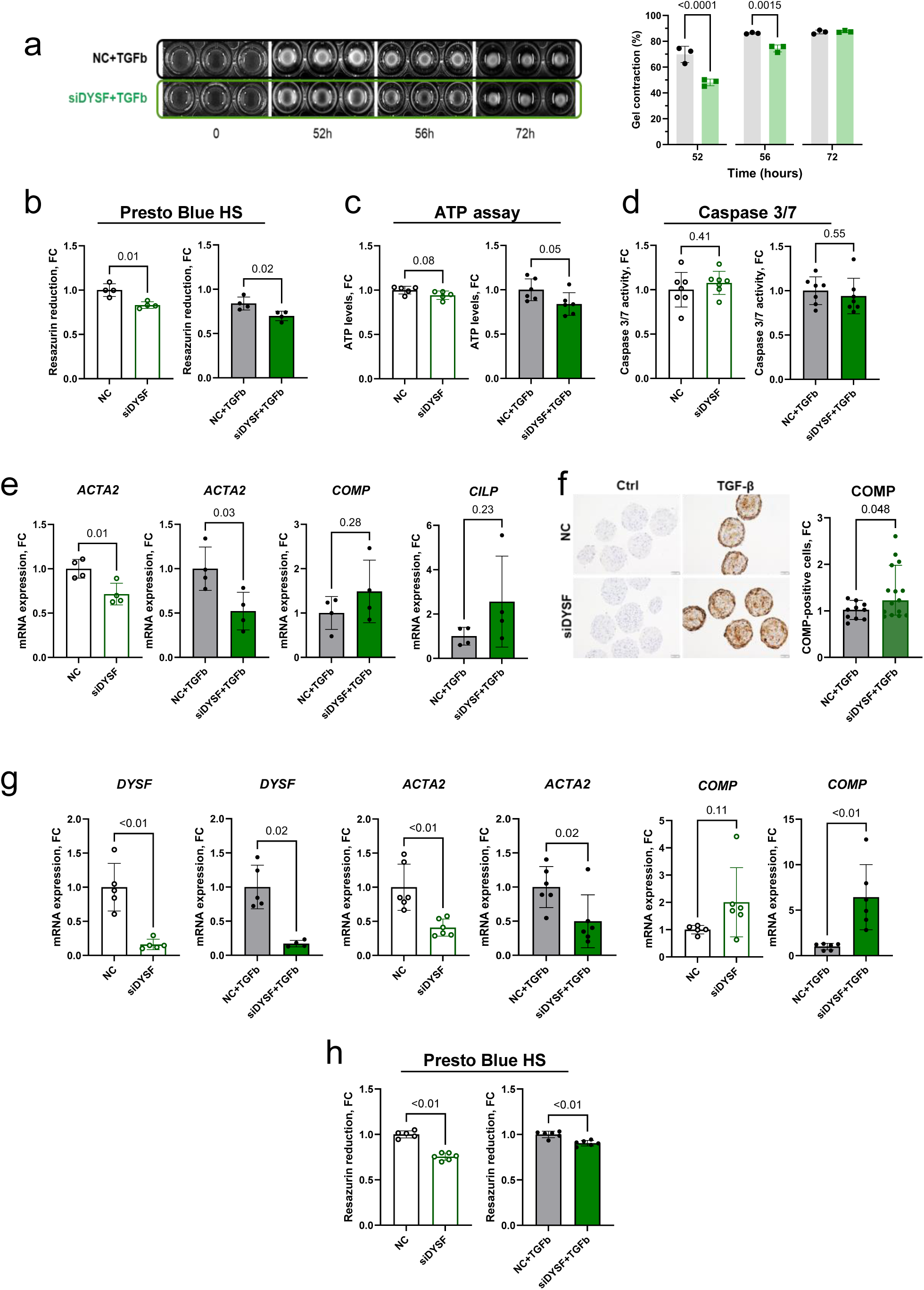
*DYSF* silencing promotes a matrifibrocyte-like phenotype in HCFs. **(a)** Contraction of 3D collagen matrices containing transfected fHCFs under profibrotic stimulation with TGF-β (20 ng/mL). The graph illustrates quantification of gel contraction based on the corresponding images (n=3; two-way ANOVA, Sidak’s multiple comparisons test; mean ± SD). **(b-d)** Effects of *DYSF* silencing on **(b)** metabolic activity (resazurin reduction), **(c)** ATP levels, and **(d)** caspase 3/7 activity in HCFs following 48 h of TGF-β stimulation (10 ng/mL; n=4-7; unpaired t-test; mean ± SD). **(e)** Relative expression of *ACTA2*, *COMP*, and *CILP* in *DYSF*-silenced fHCFs following 48 h of TGF-β stimulation (10 ng/mL), assessed by RT-qPCR (n=4; unpaired t-test; mean ± SD). **(f)** Representative images and corresponding quantification of IHC staining for COMP in cardiac microtissues (n=10-15; Mann-Whitney U test; median ± 95% CI). Nuclei were counterstained with haematoxylin. Images were acquired at 20× magnification; scale bars indicate 50 µm. **(g)** Relative mRNA expression of *DYSF, ACTA2* and *COMP* in *DYSF*-silenced adHCFs under control or TGF-β–stimulated conditions, measured by RT-qPCR (72 h; 10 ng/mL; n=4-5; unpaired t-test; mean ± SD). **(h)** Metabolic activity (resazurin reduction) in *DYSF*-silenced adHCFs after 72 h under control or TGF-β–stimulated conditions, measured using the Presto Blue HS assay (n=5-6; unpaired t-test; mean ± SD). NC: negative control; siDYSF: *DYSF* silencing.

Assessment of cellular metabolism using the PrestoBlue HS assay revealed significantly lower metabolic activity in *DYSF*-silenced fHCFs relative to NC cells under both basal and TGF-β-stimulated conditions (**Figure 3b**). Notably, no differences in metabolic activity were observed between DYSF- silenced and NC groups in iCMs or microvascular endothelial cells (ECs) (**Supplementary Figure 6**), suggesting that this effect may be specific to the fibroblast lineage. Consistent with these findings, *DYSF* silencing in fHCFs was associated with a moderate reduction in intracellular ATP levels (**Figure 3c**).

In contrast, apoptotic activity did not differ significantly between *DYSF*-silenced and NC cells under either unstimulated or TGF-β-treated conditions (**Figure 3d**). Together, these results indicate that the reductions in metabolic activity and ATP content following *DYSF* silencing are more likely attributable to impaired cellular proliferation or metabolic function rather than increased apoptosis.

The observed reductions in contractile properties and proliferative capacity of *DYSF*-silenced fibroblasts may suggest their phenotypic conversion into matrifibrocytes, a terminally differentiated, non-proliferative and non-contractile fibroblast state [23]–[25]. Supporting this hypothesis, under profibrotic conditions, *DYSF*-silenced fHCFs showed increased expression of the matrifibrocyte markers *COMP* and *CILP* compared with NC; however, these differences did not reach statistical significance (**Figure 3e**). At the same time, *DYSF*-silenced fHCFs showed reduced *ACTA2* expression compared with NC under both control and TGF-β-stimulated conditions (**Figure 3e**). To further investigate matrifibrocyte abundance in a 3D tissue context, IHC staining for COMP was performed in cardiac microtissues. Under control conditions, COMP expression was undetectable, whereas TGF-β treatment induced a marked upregulation, predominantly at the periphery of the microtissue sections, where it formed a distinct fibrotic ring (**Figure 3f**). In fibrotic microtissues, *DYSF* silencing in fHCFs resulted in a greater number of COMP-positive cells compared with NC microtissues (**Figure 3f**), suggesting an increased prevalence of a matrifibrocyte-like phenotype.

Notably, the concept of matrifibrocytes has so far been described in the context of adult human cardiac fibroblasts (adHCFs) from scarred regions of ischemic myocardium [23]. We therefore extended our investigation to adHCFs to further examine this hypothesis in a more appropriate cellular model. Consistent with fHCFs, *DYSF*-silenced adHCFs exhibited reduced *ACTA2* expression under both control and TGF-β-stimulated conditions (**Figure 3g**) and decreased metabolic activity, suggestive of reduced proliferation (**Figure 3h**). Moreover, *DYSF*-silenced adHCFs displayed increased *COMP* expression compared with NC cells, an effect that was especially pronounced under profibrotic stimulation (**Figure 3g**). Collectively, these findings indicate that *DYSF* silencing may promote cardiac fibroblast differentiation toward COMP-expressing matrifibrocytes.

### 4. COMP-positive matrifibrocytes are negative for dysferlin

To explore the abundance of the matrifibrocyte phenotype in cardiac disease, we analyzed a published CITE-seq dataset from Amrute et al.[26], which includes human left-ventricular specimens obtained from patients with acute MI (AMI; <3 months post-MI), ischemic cardiomyopathy (ICM; >3 months post-MI), and non-ischemic dilated cardiomyopathy (NICM). Our analysis confirmed the expression of matrifibrocyte markers — such as *COMP, CILP, CHAD*, and *THBS4* — in cardiac fibroblasts, which primarily co-localized within cluster FB2 (**Figure 4a-b**). Conversely, *ACTA2* expression in cardiac fibroblasts demonstrated an inverse pattern compared to the matrifibrocyte markers, showing the highest expression in clusters FB5 and FB6 (**Figure 4a, c**). Together, these observations suggest that fibroblast cluster FB2 may represent a matrifibrocyte phenotype. Furthermore, we assessed the pseudobulk expression of individual matrifibrocyte markers across the different patient groups. While all markers showed increased expression in disease, *COMP* upregulation was statistically significant across all patient groups compared to donor hearts, with the highest expression observed in the AMI cohort (**Figure 4d**). Notably, *DYSF* expression in fibroblasts was low and did not show significant differences between the conditions.

**Figure 4.**
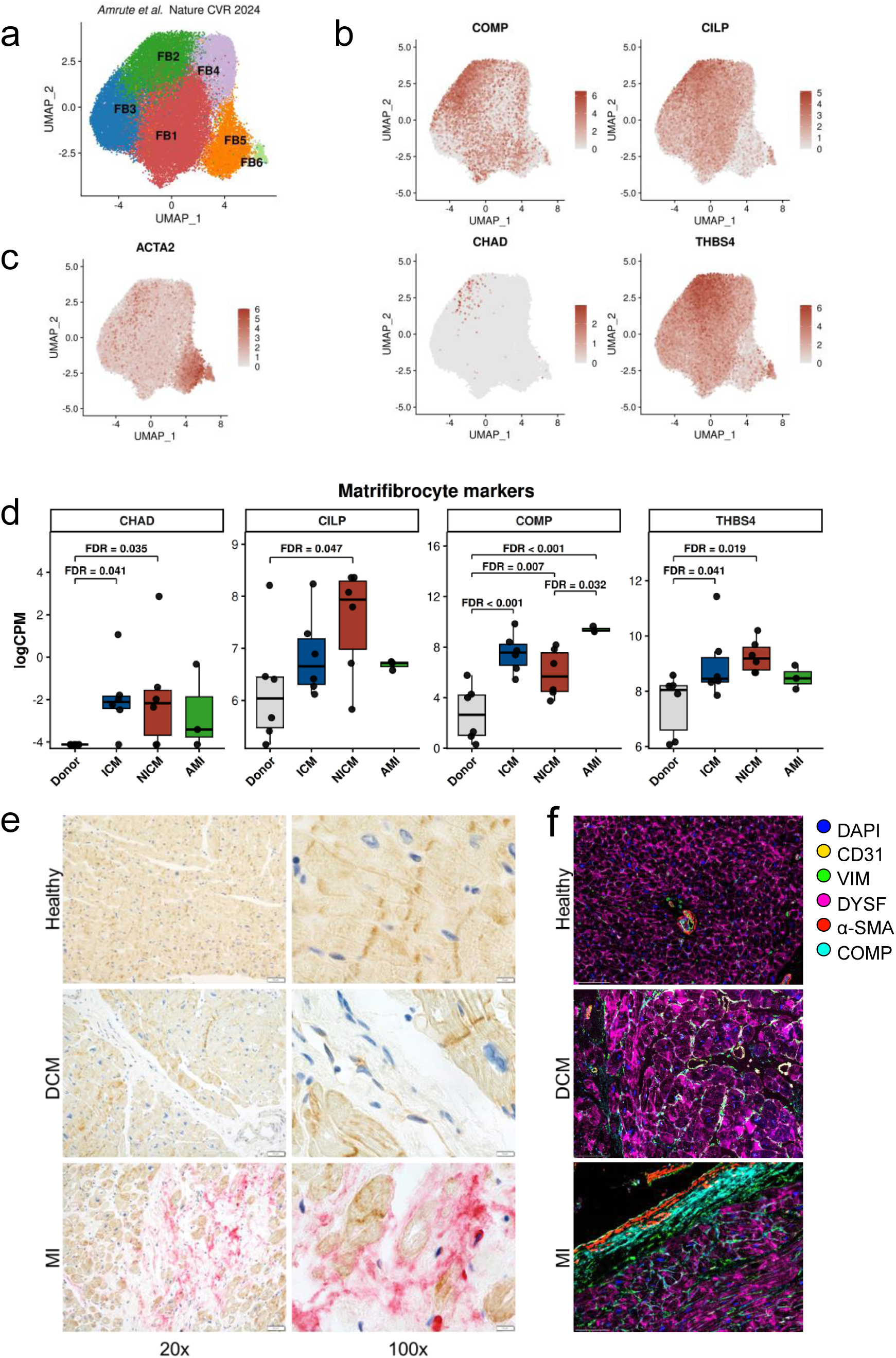
COMP-positive matrifibrocytes are negative for dysferlin. **(a)** UMAP representation of fibroblast cells from the Amrute et al. CITE-seq dataset, generated from RNA gene expression data, after removal of contamination clusters and reclustering. The dataset includes 6 donor samples and 16 heart failure samples, comprising 6 ischemic cardiomyopathy (ICM), 6 non-ischemic cardiomyopathy (NICM), and 4 acute myocardial infarction (AMI) samples. **(b)** UMAP feature plots showing expression of canonical matrifibrocyte markers *COMP*, *CILP*, *CHAD*, and *THBS4* in fibroblasts. **(c)** UMAP feature plot showing expression of the myofibroblast marker *ACTA2*. **(d)** Pseudobulk expression of matrifibrocyte markers across patient groups. Gene counts were aggregated per sample, and differential expression was tested using limma linear modeling. Contrasts included each disease group versus donor and all pairwise disease comparisons. Reported significance labels indicate Benjamini–Hochberg FDR within each significant contrast. Only samples with >100 fibroblasts were included in the differential expression analysis (Donor n = 6, ICM n = 6, NICM n = 6, AMI n = 3). **(e)** Representative images of COMP (red) and dysferlin (brown) IHC in healthy myocardium, dilated cardiomyopathy (DCM) and myocardial infarction (MI) tissue sections. Nuclei were counterstained with haematoxylin. **(f)** Representative immunofluorescence images acquired using the COMET platform (Lunaphore). Nuclei were counterstained with DAPI. Scale bars: 100 µm.

To validate these results, we performed IHC staining on myocardial tissue sections from healthy, DCM, (NICM), and MI samples, which confirmed a significant abundance of COMP in MI tissue (**Figure 4e**). Importantly, COMP-positive fibroblasts, which were highly abundant in MI tissue, were negative for dysferlin. In contrast, DCM tissue was enriched in dysferlin-positive/COMP-negative fibroblasts, further supporting a regulatory relationship between dysferlin and COMP expression. Additionally, we performed multiplex immunofluorescence (IF) staining of human myocardial tissue sections using the Lunaphore COMET platform. The staining confirmed the increased abundance of the classical fibrotic marker periostin in both DCM and MI myocardium, whereas α-SMA expression was largely confined to the vascular walls (**Figure 4f, Supplementary Figure 7**). As observed in IHC, expression of the matrifibrocyte-associated marker COMP was detected in MI. This observation supports a role for COMP-producing matrifibrocytes in shaping the pathological tissue architecture of the HF myocardium. Dysferlin expression was predominantly observed in CMs but was also present in HCFs specifically in DCM (**Supplementary Figure 7**). Notably, dysferlin-expressing HCFs in fibrotic regions were COMP-negative (**Supplementary Figure 7**). In contrast, in healthy control myocardium dysferlin expression was restricted to CMs.

### 5. Dysferlin is involved in the regulation of the TGF-β-FOSL2-autophagy axis

Fibroblast activation and profibrotic differentiation, accompanied by enhanced ECM synthesis, are known to be mediated by the AP-1 transcription factor FOSL2, which acts downstream of TGF-β signalling [5], [7]. Given that *DYSF* silencing increased the expression of ECM-related markers and promoted HCF differentiation into matrifibrocytes, we hypothesized that this effect might involve FOSL2 regulation. Accordingly, *DYSF*-silenced fHCFs showed increased protein levels of FOSL2 under both control and TGF-β-stimulated conditions (**Figure 5a**). In turn, *FOSL2* silencing significantly increased dysferlin levels in TGF-β-stimulated fHCFs (**Figure 5b**), indicating reciprocal regulation between FOSL2 and dysferlin. Based on this observation, we further hypothesized that increased FOSL2 expression might suppress dysferlin levels in cardiac fibroblasts. To explore this regulatory relationship in an *in vivo* setting, we used transgenic mice overexpressing Fosl2 (Fosl2^tg^) [5], [8]. Consistent with our hypothesis, we observed decreased dysferlin levels in Fosl2-overexpressing cardiac fibroblasts (**Figure 5c**). Notably, the downregulation of dysferlin in Fosl2^tg^ fibroblasts was detected at the protein level, whereas *Dysf* mRNA expression did not differ significantly, confirming lack of correlation between dysferlin transcript levels and protein abundance (**Figure 5d**).

**Figure 5.**
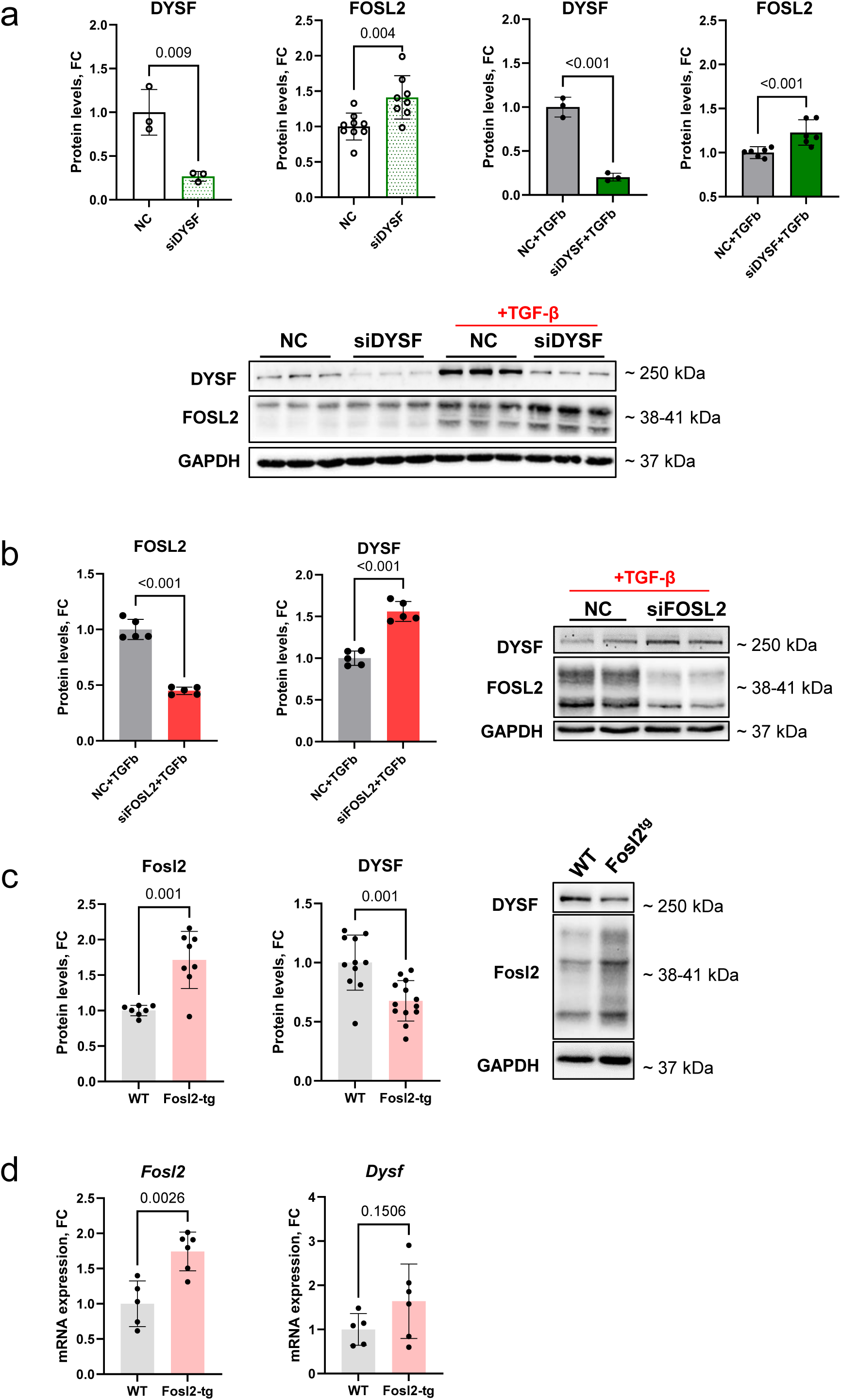
Reciprocal regulation between dysferlin and FOSL2 in cardiac fibroblasts. **(a)** Representative immunoblots and quantification of FOSL2 and dysferlin (DYSF) protein levels in *DYSF*-silenced fHCFs following 72 h under control or TGF-β–stimulated conditions (n=5; unpaired t-test; mean ± SD). **(b)** Quantification of DYSF and FOSL2 protein levels in *FOSL2*-silenced fHCFs with corresponding representative immunoblots. **(c)** Representative immunoblots and quantification of Fosl2 and dysferlin protein levels in cardiac fibroblasts from wild-type (WT) and Fosl2^tg^ mice (n=5-13; unpaired t-test; mean ± SD). **(d)** Relative gene expression of *Fosl2* and *Dysf* in WT and Fosl2^tg^ cardiac fibroblasts, measured by RT-qPCR (n=5-6; unpaired t-test; mean ± SD).

Remarkably, FOSL2 has previously been shown to regulate autophagy in cardiac fibroblasts, and the TGF-β–FOSL2–autophagy axis has been implicated in myofibroblast differentiation [7]. Here, *DYSF* silencing upregulated the key autophagy marker microtubule-associated protein 1 light chain 3 (LC3BII), while downregulating the autophagy receptor P62 in TGF-β-stimulated fHCFs (**Figure 6**), suggesting induction of autophagy either directly or via FOSL2 upregulation. Together, these results indicate that the profibrotic effects of *DYSF* silencing in cardiac fibroblasts may involve the TGF-β–FOSL2– autophagy axis.

**Figure 6.**
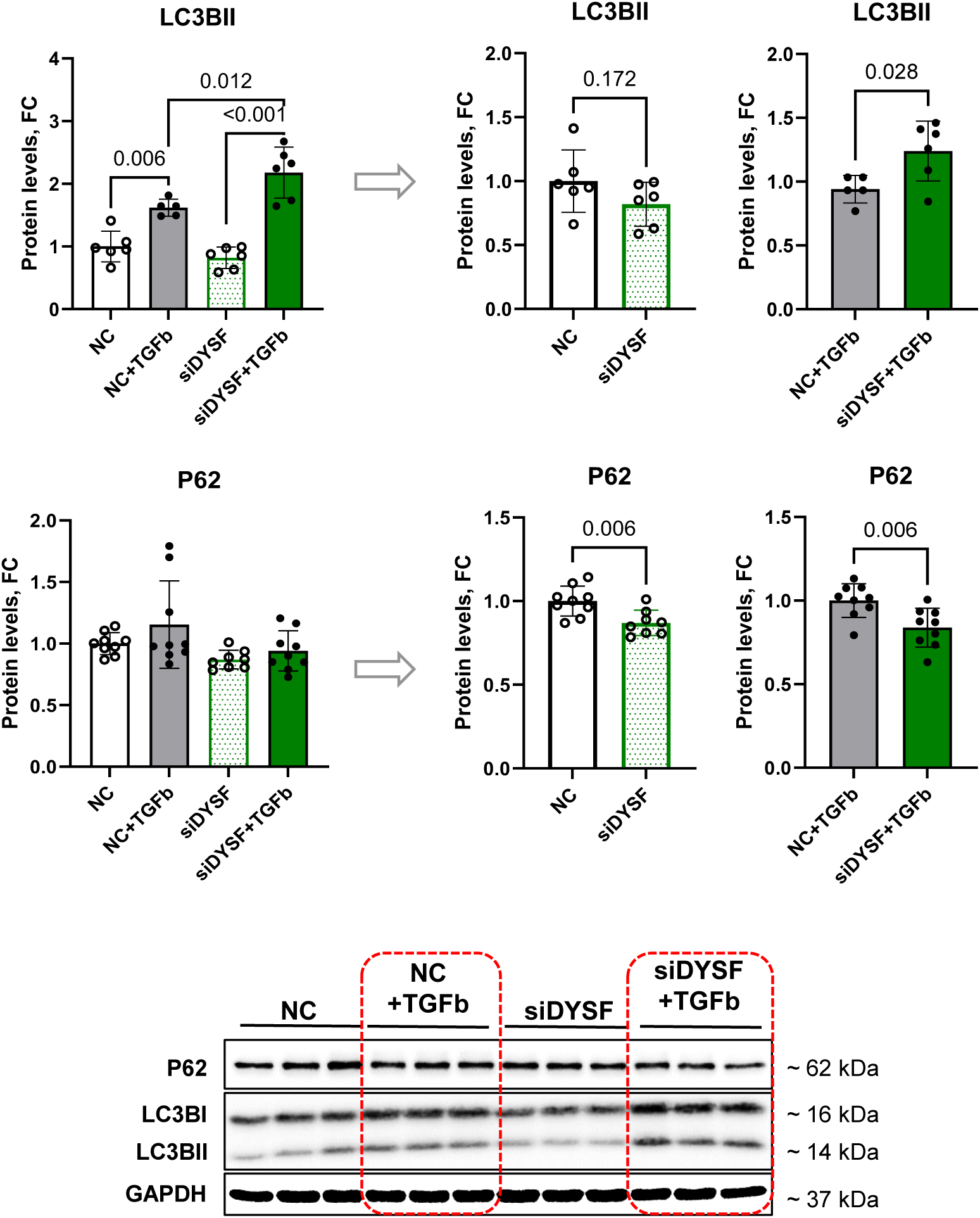
Dysferlin involvement in regulating autophagy-related proteins. Quantification of the autophagy-related proteins LC3BII and P62 with corresponding representative immunoblots. Analyses were performed in transfected fHCFs following 72 h under control or TGF-β-stimulated conditions (n=6-9; one-way ANOVA with Tukey’s multiple comparisons test for four experimental conditions or unpaired t-test for two experimental conditions; mean ± SD). NC: negative control; siDYSF: *DYSF* silencing.

### 6. Dysferlin expression in HCFs is regulated by MXRA5

Among other fibrotic markers, *MXRA5* emerges as one of the most consistently upregulated genes in the HF transcriptome of total myocardium according to the Reference of the HF Transcriptome (ReHeaT; **Supplementary Figure 5**). To validate this observation specifically in cardiac fibroblasts, we performed pseudobulk differential expression analysis of published snRNA-seq datasets from human DCM myocardium (GSE183852 [14], EGAS00001006374 [16], SCP1303 [15], GSE217494 [26]). The analysis revealed significant upregulation of *MXRA5* expression in DCM fibroblasts, which was consistent across all datasets (**Figure 7a**).

**Figure 7.**
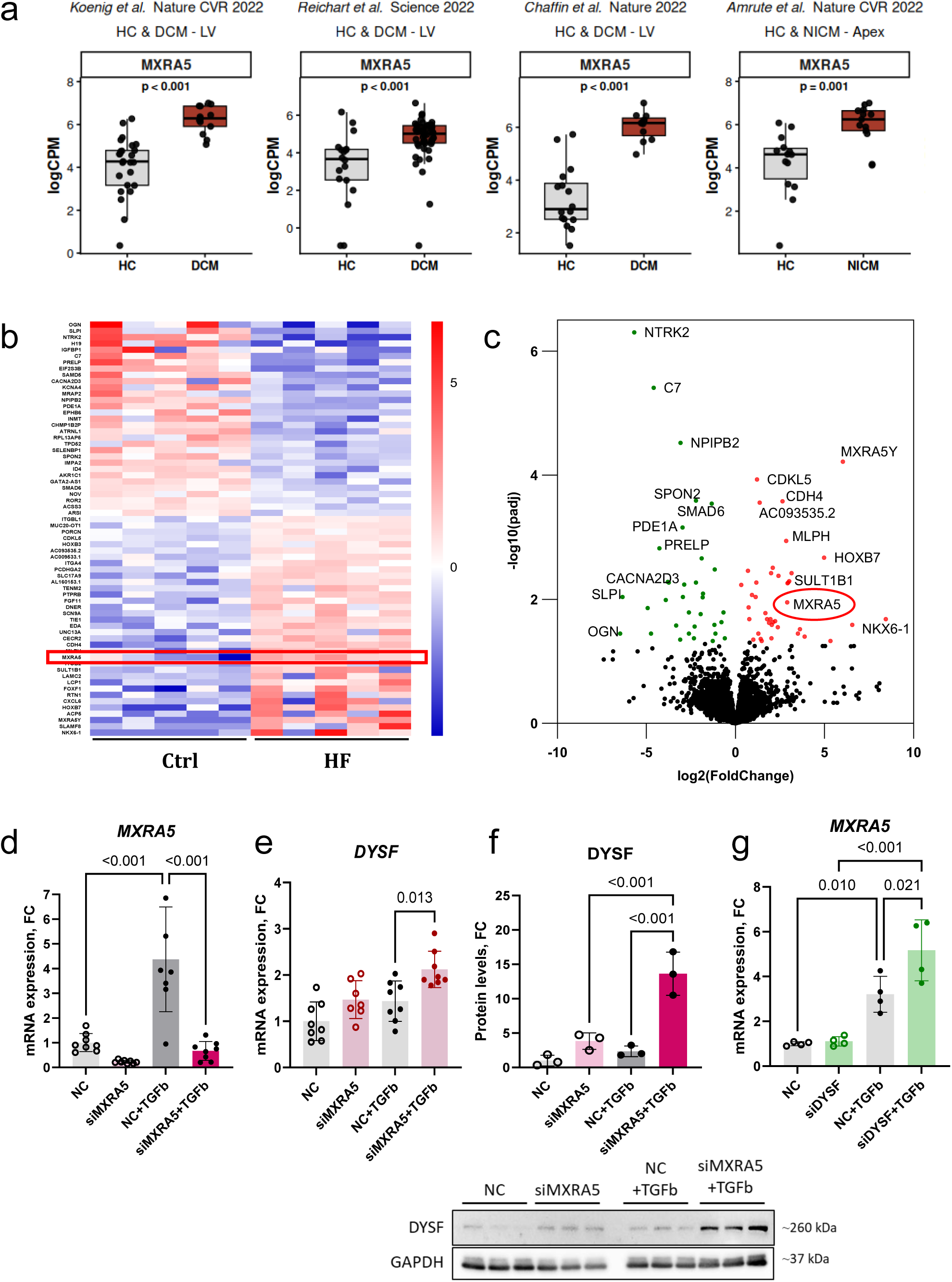
Dysferlin expression in HCFs is regulated by MXRA5. **(a)** Pseudobulk expression of MXRA5 in the datasets from Koenig et al. (GSE183852), Reichart et al. (EGAS00001006374), Chaffin et al. (SCP1303) and Amrute et al. (GSE217494). HC: healthy control; DCM: dilated cardiomyopathy; NICM: non-ischemic cardiomyopathy. **(b)** Heatmap and **(c)** volcano plot visualising global transcriptomic changes in human atrial cardiac fibroblasts from HF patients and healthy myocardium. Bulk RNA sequencing was performed using the Illumina HiSeq platform, and differentially expressed genes were identified using DESeq2 (n=5; absolute log₂FC > 1; adj. p < 0.05). *MXRA5* is highlighted in red. **(d)** *MXRA5* silencing in fHCFs and its effect on **(e)** *DYSF* gene expression following 48 h of TGF-β stimulation (10 ng/mL, n=7-8; one-way ANOVA, Tukey’s multiple comparisons test; mean ± SD). **(f)** Quantification of dysferlin protein levels in *MXRA5*-silenced fHCFs 72 h after TGF-β stimulation and corresponding immunoblots (n=3; one-way ANOVA, Tukey’s multiple comparisons test; mean ± SD). **(g)** Relative expression of MXRA5 in transfected fHCFs following 48 h of TGF-β stimulation (10 ng/mL) (n = 4; unpaired t-test; mean ± SD). NC: negative control; siMXRA5: *MXRA5* silencing; siDYSF: *DYSF* silencing.

In line with these findings, our bulk RNA sequencing of HCFs isolated from patients with end-stage HF (n = 5) and unaffected donor hearts (n = 5) identified *MXRA5* among the significantly upregulated genes in HF (log2FC = 2.91, adjusted p < 0.05) (**Figure 7b-c**). We therefore next sought to investigate whether MXRA5 regulates DYSF expression by silencing *MXRA5* in fHCFs. Our results demonstrated that *MXRA5* silencing in fHCFs led to upregulation of dysferlin expression at both the mRNA and protein levels (**Figure 7d-f**). Interestingly, *DYSF* silencing in fHCFs was associated with increased *MXRA5* expression under fibrotic conditions (**Figure 7g**), suggesting a potential reciprocal regulatory relationship between MXRA5 and dysferlin.

Overall, our results point to a regulatory role for MXRA5 in controlling dysferlin expression in fHCFs.

### 7. Dysferlin may undergo post-translational modifications

Notably, our proteomic and transcriptomic analyses of HCFs from HF patients and donor hearts yielded two distinct sets of differentially expressed candidates with no overlap (**Figure 1b-c**, **Figure 7b-c**). While dysferlin emerged as the most upregulated protein in HF-HCFs, transcriptomic analysis did not identify *DYSF* among the significantly deregulated genes. Furthermore, analysis of public sc/snRNA-seq datasets identified CMs, as the primary producers of dysferlin within myocardial tissue, both in healthy (ERP123138 [27]) and HF conditions (SCP1303 [15], GSE183852 [14], EGAS00001006374 [16]) (**Supplementary Figure 8a**). Differential gene expression analysis across individual datasets yielded inconsistent results with two of the three datasets showing downregulation of *DYSF* in DCM fibroblasts **(Supplementary Figure 8b-d**), contrasting with the results obtained in our proteomic analysis. A discrepancy between dysferlin mRNA and protein levels was also observed in Fosl2^tg^ mouse cardiac fibroblasts, where downregulation of dysferlin protein, compared with cardiac fibroblasts from WT mice, was not accompanied by corresponding changes in *Dysf* mRNA expression (**Figure 5c-d**). Consistent with these observations, data retrieved from the Open Targets Platform indicated a low correlation between dysferlin mRNA and protein expression across different organs, including the heart (**Supplementary Figure 9**).

These observations suggest, among other possibilities, the involvement of post-translational modifications in regulating dysferlin protein stability. In particular, S-acylation and its most common form, S-palmitoylation, has been reported to protect membrane proteins from degradation by enhancing their association with the cell membrane [28]. Consistent with this, analysis of palmitoylome datasets via the SwissPalm database identified dysferlin as a predicted S-acylated protein, including in human myocardial tissue [29] (**Supplementary Figure 10a**), and sequence analysis revealed multiple candidate cysteine sites, including high-confidence predictions by CSS-Palm 4.0 (**Supplementary Figure 10b-c**). In addition, “protein palmitoylation” (GO:0045234) was enriched in HF cardiac fibroblasts (**Figure 8a**). Pharmacological inhibition of S-acylation with 2-bromopalmitate (2-BP) reduced dysferlin levels in iCMs and ECs (**Figure 8b-c**), paralleling the response of the S-acylated protein flotillin-1.

**Figure 8.**
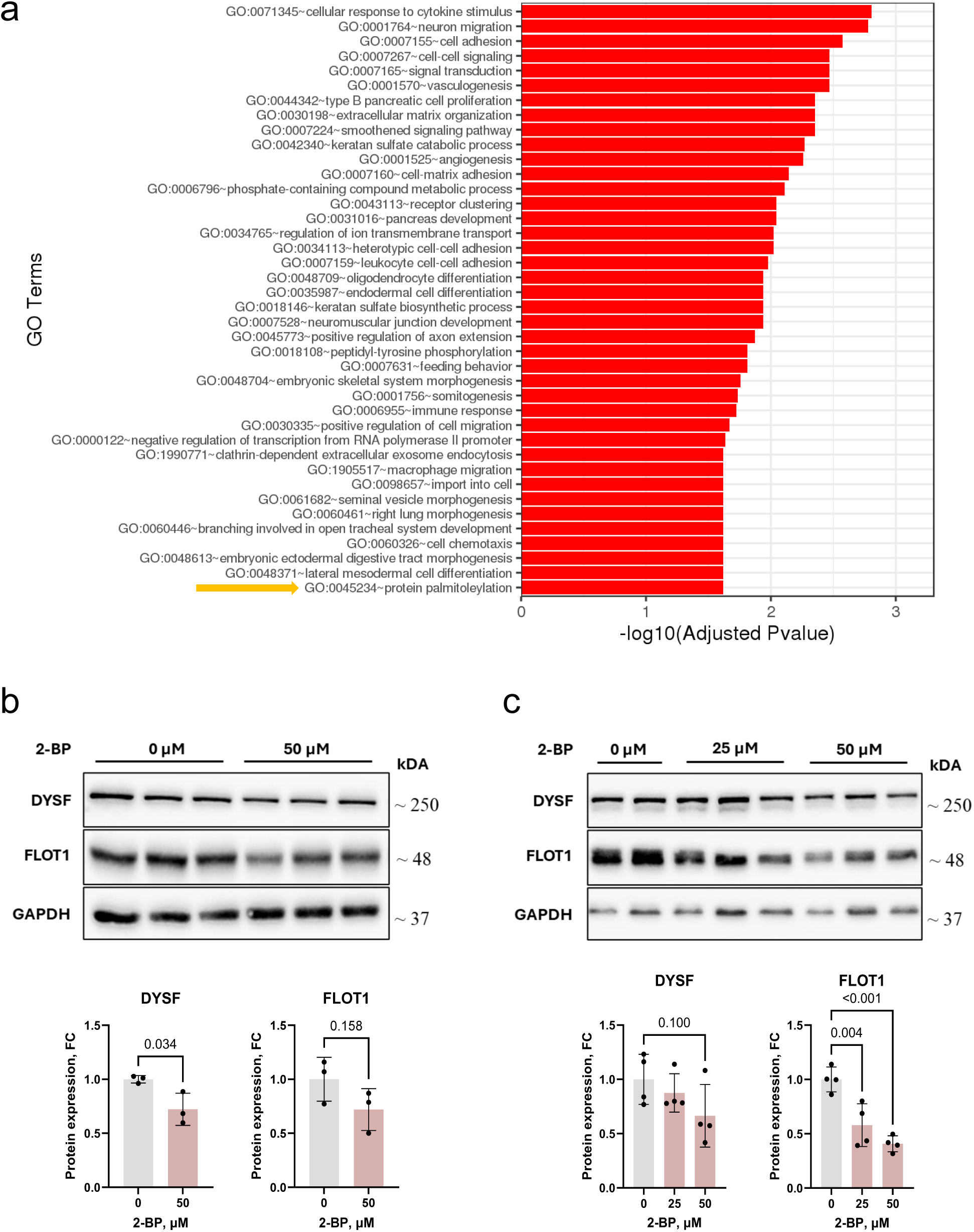
Dysferlin may undergo post-translational modifications. **(a)** Gene Ontology enrichment in HF HCFs. Significantly deregulated genes were grouped by Gene Ontology (GO) terms, and enrichment was tested using Fisher’s exact test (GeneSCF v1.1-p2). The plot displays GO terms that are significantly enriched (adj. p-value < 0.05) in the differentially expressed gene sets. **(b)** Dysferlin and flotillin-1 protein levels in iCMs following 24 h of treatment with 2-bromopalmitate (2-BP). **(c)** Dysferlin and flotillin-1 protein levels in ECs following 24 h of treatment with 2-BP.

Together, these data suggest that dysferlin is likely S-acylated, which may contribute to the discrepancy between dysferlin mRNA and protein expression levels.

## Discussion

We identify dysferlin as a stress-induced antifibrotic regulator that limits fibroblast differentiation toward a COMP-positive matrifibrocyte state in human HF. Most of our current understanding of dysferlin function arises from studies on dysferlinopathies, a group of disorders caused by mutations in the *DYSF* gene that lead to progressive muscular dystrophy [30]. The pathological phenotype in dysferlinopathies primarily affects skeletal muscle, whereas the prevalence and extent of cardiac involvement remain subjects of debate [31]. In mouse models of dysferlinopathy, dysferlin deficiency has been shown to induce exercise-induced cardiomyopathy characterized by chamber dilation and systolic dysfunction [20]. Furthermore, while substantial evidence supports a cytoprotective role of dysferlin in muscle cells [32], its function in fibroblasts has not been extensively investigated.

Previous studies have shown that murine dysferlin-deficient fibroblasts, similar to dysferlin-deficient myoblasts, exhibit defects in intracellular vesicular trafficking and membrane fusion processes. [33]. Dysferlin has also been detected in skin fibroblasts, where its reduced protein levels are associated with impaired membrane resealing and defective blebbing under hypotonic shock *in vitro* [34]. However, how these observations translate to fibroblast function remains unclear. Notably, experimental improvement of membrane repair defects in dysferlin-deficient mice failed to prevent progressive dystrophic changes in muscle tissue, suggesting that dysferlin function is not limited solely to membrane repair [35].

Supporting a broader biological role for dysferlin, our study is the first to demonstrate its involvement in the regulation of cardiac fibroblast activation. Our data demonstrate that *DYSF* silencing in HCFs increased the expression of ECM-related markers and promoted a more pronounced fibrotic phenotype in engineered cardiac microtissues. Notably, dysferlinopathies have been linked to fibrotic changes in skeletal muscle, including increased collagen deposition within muscle tissue [36], which has largely been considered a consequence of progressive muscle damage [37], [38]. In this regard, our observations provide novel insight into the aetiology of fibrosis in dysferlinopathies, suggesting that dysferlin deficiency itself may promote fibroblast activation and enhanced collagen synthesis, thereby contributing to the development of fibrotic phenotype.

In addition to enhanced ECM production, *DYSF* silencing in HCFs promoted a phenotypic transition toward matrifibrocytes, a terminally differentiated state characterized by a quiescent, non-proliferative phenotype. The concept of matrifibrocytes was first described by Fu and colleagues using lineage tracing in a murine myocardial infarction model, with subsequent identification of COMP-positive matrifibrocytes in human infarcted myocardium [23]. More recent studies have employed the term matrifibrocytes or matrifibrocyte-resembling cells to describe a matrix-producing, α-SMA-negative fibroblast population expressing chondrocyte- and osteoblast-associated markers [24]–[26]. Due to their distinct secretory profile, including upregulation of COMP, CILP, CHAD, and THBS4, matrifibrocytes contribute to scar maturation and long-term maintenance of the post-injury myocardial microenvironment, thereby supporting scar integrity and preventing ventricular wall rupture after myocardial infarction [39]. In the present study, IHC and IF analyses confirmed robust COMP expression in infarcted myocardium. Importantly, COMP-positive matrifibrocytes lacked dysferlin expression, whereas dysferlin-positive HCFs were COMP-negative, corroborating our *in vitro* findings and supporting an inverse relationship between dysferlin expression and the matrifibrocyte phenotype. Collectively, these findings suggest that dysferlin acts as an endogenous negative regulator of COMP-positive matrifibrocyte differentiation.

Mechanistically, profibrotic differentiation of *DYSF*-silenced HCFs was accompanied by upregulation of the transcription factor FOSL2, a recognized driver of pathological extracellular matrix remodelling and myocardial fibrogenesis [5], [7], [40]. Conversely, silencing of *FOSL2* resulted in increased dysferlin protein levels, suggesting a reciprocal regulatory relationship between FOSL2 and dysferlin. Consistent with these findings, cardiac fibroblasts derived from Fosl2-overexpressing mice exhibited reduced dysferlin levels, further supporting the bidirectional regulation.

Notably, *MXRA5* silencing increased dysferlin expression in HCFs, suggesting that both FOSL2 and MXRA5 act as TGF-β–induced negative regulators of dysferlin in cardiac fibroblasts. MXRA5 has been implicated in pathological ECM remodelling [41] and emerged as one of the most consistently upregulated genes in HF in our analysis of published transcriptomic datasets, corroborating previous transcriptomic analyses [42], [43]. Despite its upregulation in HF, the precise functional role of MXRA5 in cardiac fibrogenesis remains unclear. In this context, our findings identify dysferlin as a potential downstream target of MXRA5 in cardiac fibroblasts.

In addition to upregulated FOSL2 expression, *DYSF* silencing in HCFs was associated with differential regulation of the autophagy-related proteins LC3BII and P62, which may reflect increased autophagic flux. This observation aligns with a previous study reporting elevated autophagy in Fosl2-overexpressing mouse heart fibroblasts [7]. Moreover, previous reports of increased accumulation of LAMP2-positive lysosomes in dysferlin-deficient fibroblasts [33] further support a role for dysferlin in the regulation of the autophagy–lysosomal pathway. In the heart, autophagy is involved in a wide range of biological processes and also exerts cardioprotective effects [13]. However, enhanced autophagy in cardiac fibroblasts promotes their phenotypic conversion into an activated state and is consistently associated with fibrotic remodelling of myocardial tissue [7]. Moreover, TGF-β-induced fibrosis has been shown to depend on autophagy induction [44]. Consequently, the ability of dysferlin to act as a negative regulator of the TGF-β-FOSL2-autophagy axis in HCFs suggests its potential antifibrotic role. The observed upregulation of dysferlin mediated by TGF-β signalling may represent an endogenous compensatory mechanism in response to myocardial injury. Such endogenous pro-regenerative responses in the adult heart have already been reported in fibrotic mice hearts after MI [45], [46]. In this light, supporting endogenous pro-regenerative mechanisms within myocardial tissue emerges as a potential therapeutic strategy to attenuate cardiac fibrosis. Studies on dysferlinopathy provide a solid foundation for strategies to increase dysferlin expression, including intramuscular or systemic vector delivery [37], [47]–[49]. Importantly, experimental *DYSF* overexpression in a non-human primate model showed no significant side effects or safety concerns [47]. Moreover, dysferlin is known to exert cytoprotective effects in cardiomyocytes by maintaining membrane integrity and proper organization of the tubular system under stress conditions [50]–[53]. Therefore, our findings, together with previously published reports, demonstrate the pro-regenerative potential of dysferlin in myocardial tissue, positioning it as a potential therapeutic candidate for counteracting adverse cardiac remodelling.

Importantly, our study demonstrated a poor correlation between dysferlin mRNA expression and protein abundance, potentially due to post-translational modifications (PTMs). Publicly available databases identified dysferlin as a potentially S-acylated protein. S-acylation enhances membrane association and stability of membrane proteins by increasing their hydrophobicity and protecting them from degradation [28], [54], [55]. Accordingly, we hypothesize that S-acylation may stabilise dysferlin at the plasma membrane and thereby reduce its degradation. Although our experiments with inhibition of S-acylation support this possibility, further in vitro validation is required to confirm dysferlin S-acylation and its functional implications.

In conclusion, the present study places dysferlin into a fundamentally new context of cardiac fibroblast biology and, for the first time, suggests that dysferlin restrains fibroblast differentiation into COMP-positive matrifibrocytes. We demonstrate that dysferlin acts as a negative regulator of the TGF-β– FOSL2–autophagy axis, with both FOSL2 and MXRA5 functioning as upstream TGF-β-induced repressors of dysferlin expression. This TGF-β-driven upregulation of dysferlin reflects an endogenous compensatory mechanism that restrains myocardial fibrogenesis. Collectively, these findings position dysferlin as a novel regulator of cardiac fibroblast fate in heart failure and a potential target for limiting adverse myocardial remodelling.

## Acknowledgements

We thank Francesca Barone (Department of Clinical Pharmacology and Toxicology, University Hospital Zurich) for her contribution to the S-acylation experiments. We acknowledge the Functional Genomics Center Zurich for proteomics analysis, Genewiz (Leipzig, Germany) for bulk transcriptomics analysis, the iPSC Core Facility (Institute for Regenerative Medicine, University of Zurich) for providing differentiated iCMs, and the Center for Microscopy and Image Analysis (University of Zurich) for support with the Lunaphore COMET multiplex platform. We also thank Nick Li and Pal Johansen (Department of Dermatology, University Hospital Zurich) for their technical assistance.

## Author contributions

I.K. and G.K. contributed to the conception and design of the work. I.K., M.G., A.L., L.M., D.N., F.D.M., G.A.B., E.P., F.R., M.M., P.L., M.V., O.D., P.B., and G.K. generated, analysed, or interpreted the data. I.K. and G.K. wrote the manuscript. M.V., M.M. and P.B. helped edit the manuscript. G.K. funded the study. All authors read and approved the final manuscript.

---

**Supplementary Figure 1.**
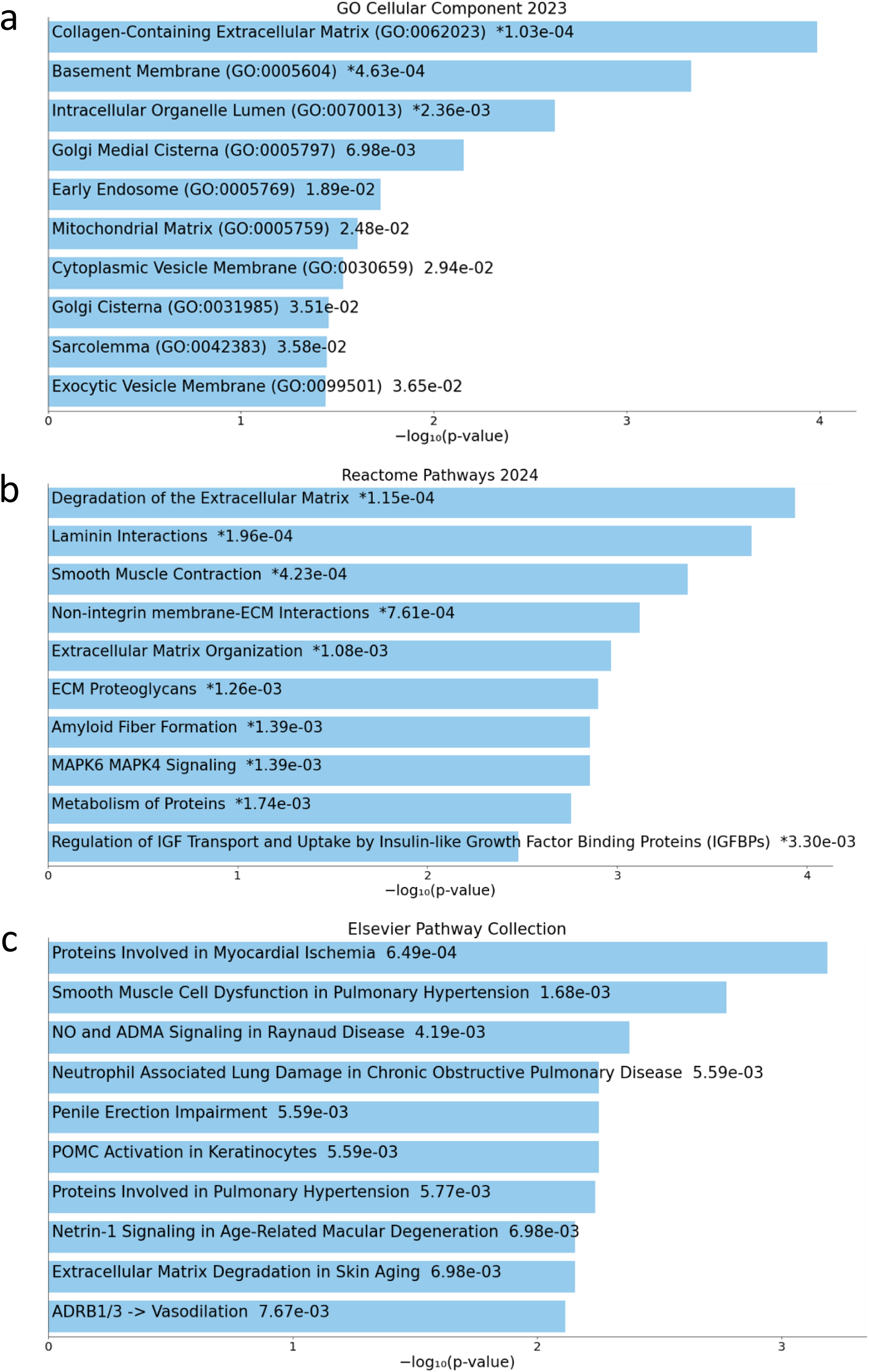
Enrichment analysis of differentially expressed proteins in HF fibroblasts. Bar charts display the top enriched terms from selected EnrichR libraries, along with the corresponding p-values (< 0.05): **(a)** GO Cellular component 2023, **(b)** Reactome pathways 2024, **(c)** Elsevier Pathway Collection. An asterisk (*) indicates terms with a significant adjusted p-value (< 0.05). Visualised using the Enrichr Appyter.

**Supplementary Figure 2.**
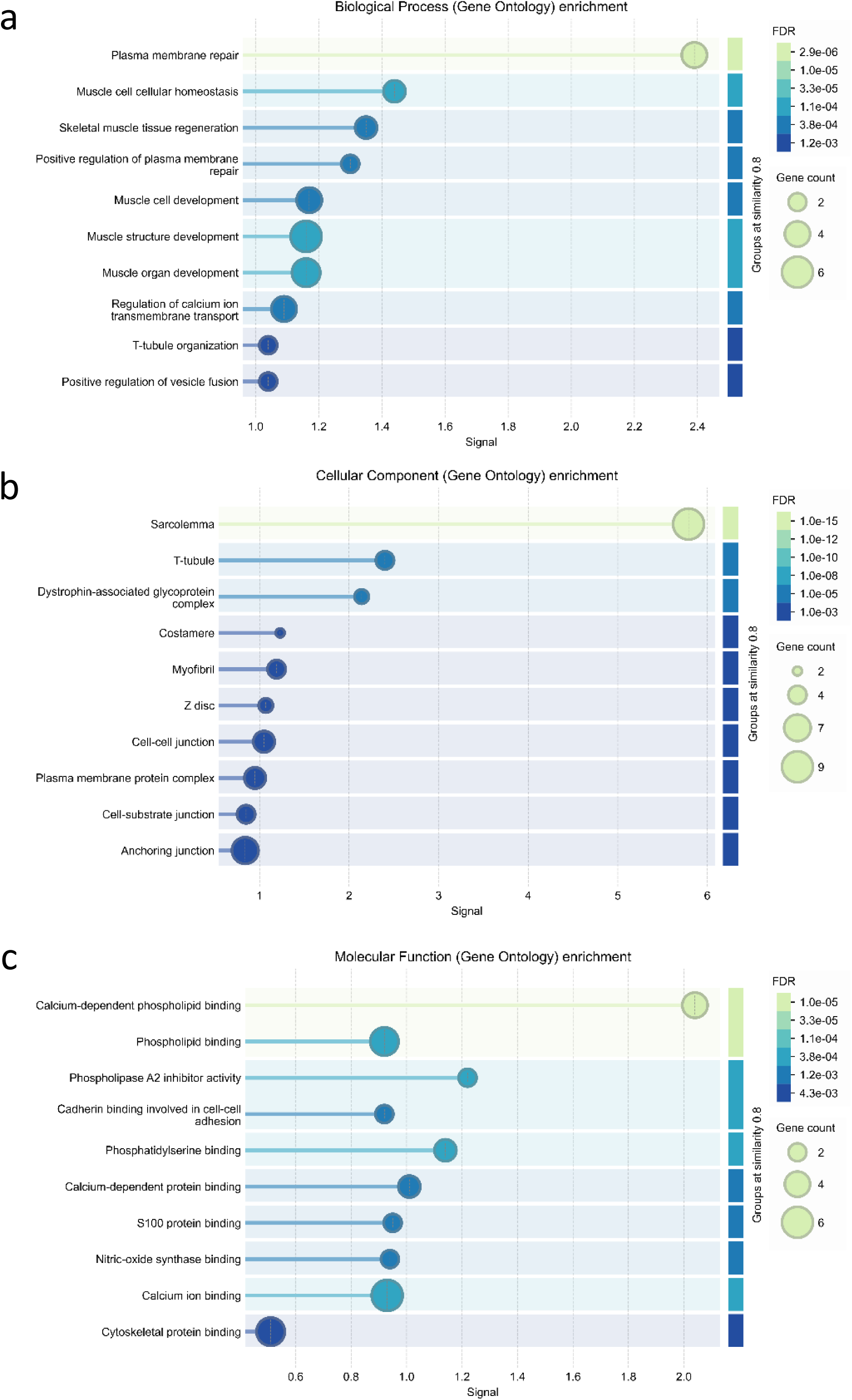
Functional enrichment visualisation for dysferlin. Charts display the top enriched GO terms for **(a)** Biological Process, **(b)** Cellular Component and **(c)** Molecular Function. Data were retrieved from STRING (https://string-db.org/, assessed on 11.12.25).

**Supplementary Figure 3.**
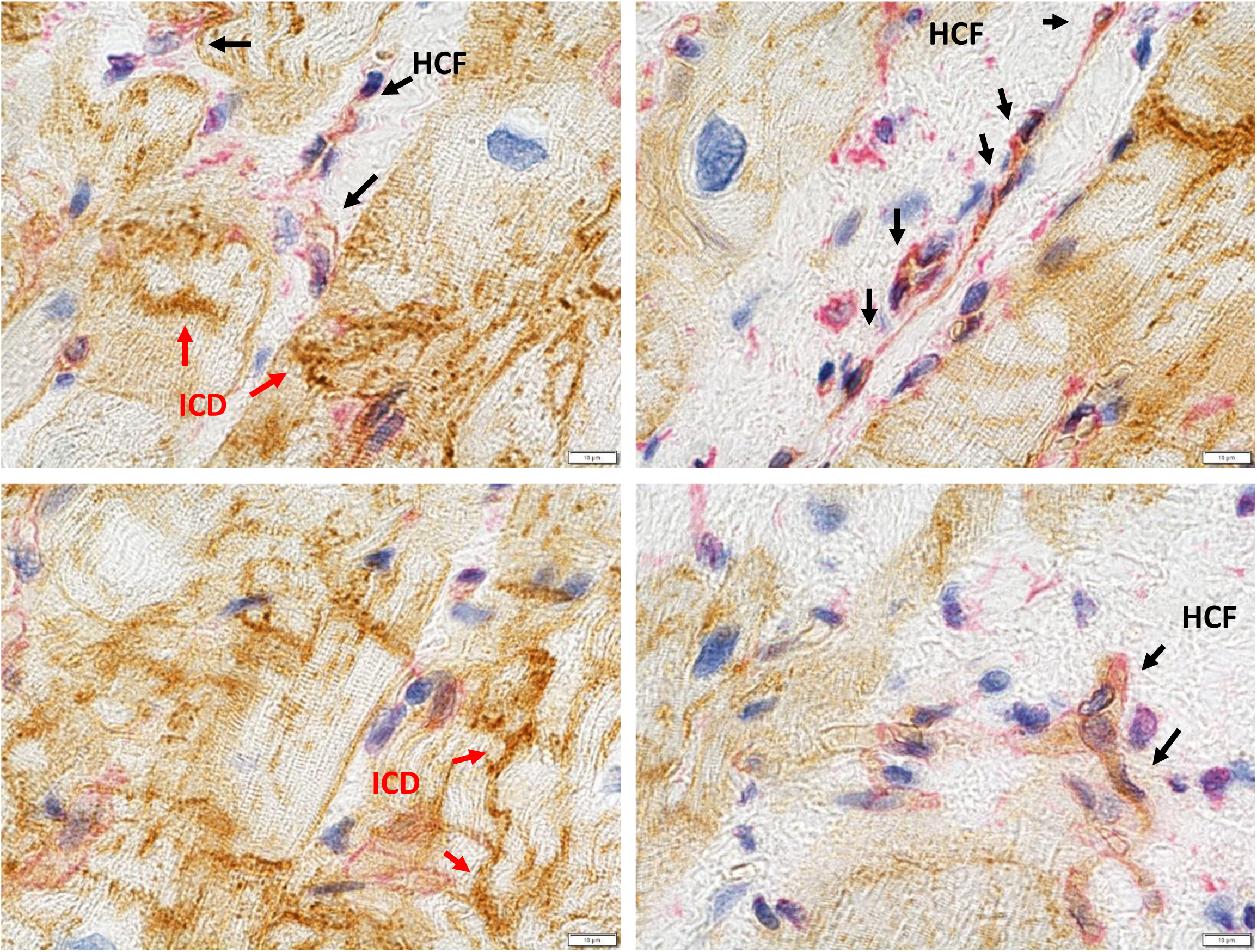
Representative images of IHC staining for dysferlin and vimentin in left ventricular tissue sections from DCM myocardium. Arrows indicate HCFs and intercalated discs (ICD). Scale bars: 10 µm.

**Supplementary Figure 4.**
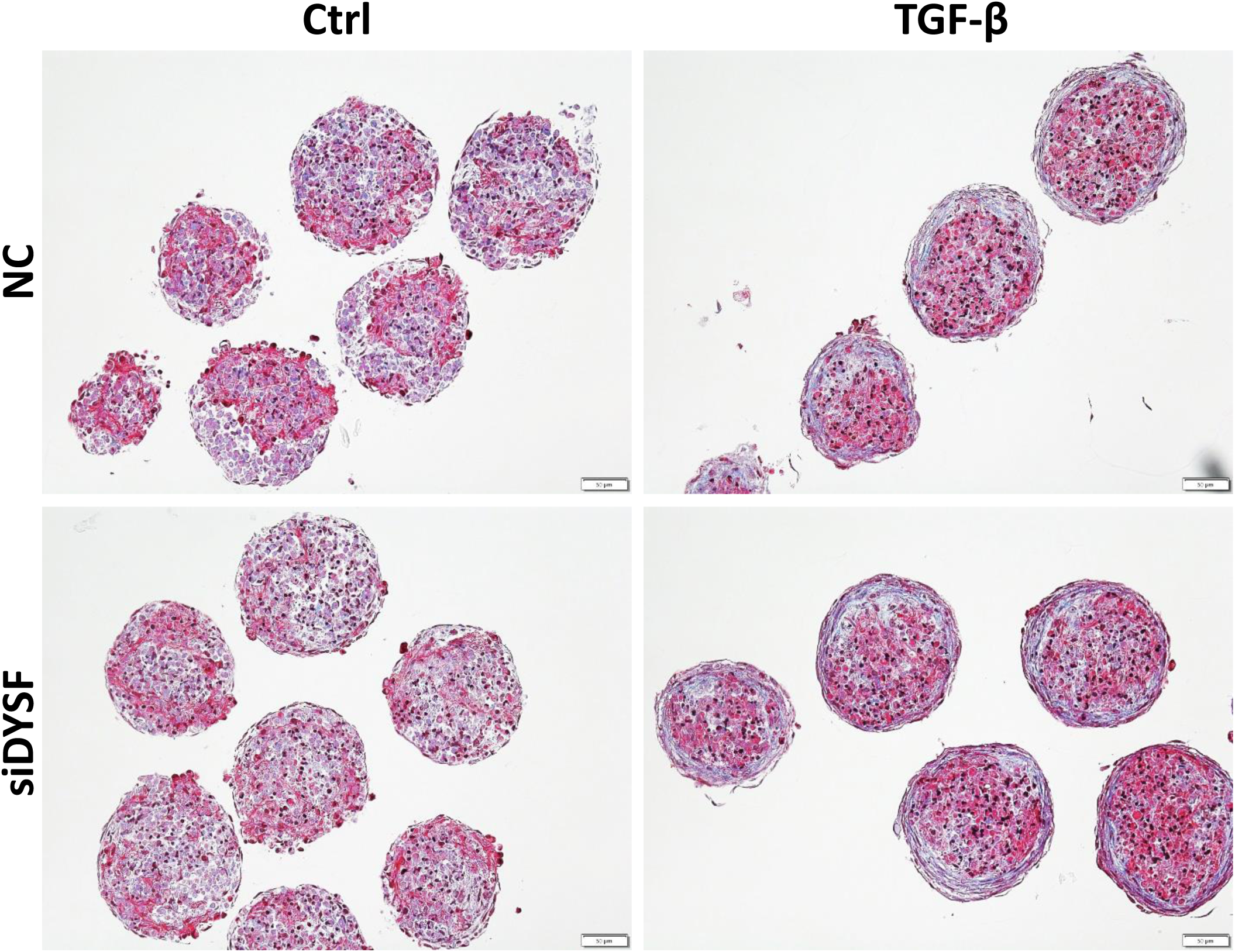
Masson’s trichrome staining of cardiac microtissues following 6 days of TGF-β stimulation. Scale bar: 50 μm.

**Supplementary Figure 5.**
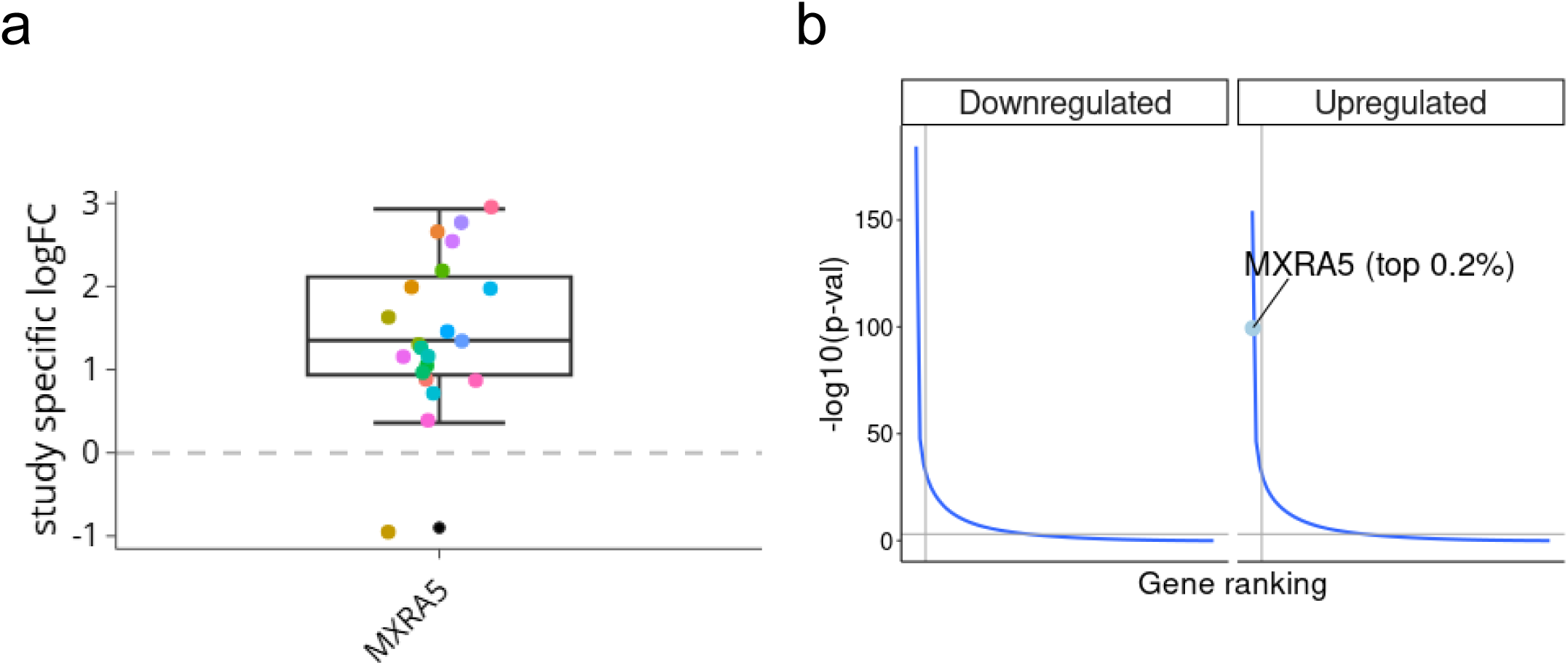
*MXRA5* as a top-regulated gene in HF transcriptome. **(a)** Comparison of the reported log2 fold change for *MXRA5* expression across 20 individual bulk transcriptomic studies on end-stage HF that were integrated into the HF consensus signature meta-analysis by Lanzer et al. (2025). **(b)** The position of *MXRA5* within the consensus HF gene ranking derived from the meta-analysis using Fisher’s combined p-value. The vertical line marks the top 500 genes; the horizontal line shows a Fisher p-value threshold of 0.05. The graphs were generated using the ReHeat2-App (https://saezlab.shinyapps.io/reheat2/).

**Supplementary Figure 6.**
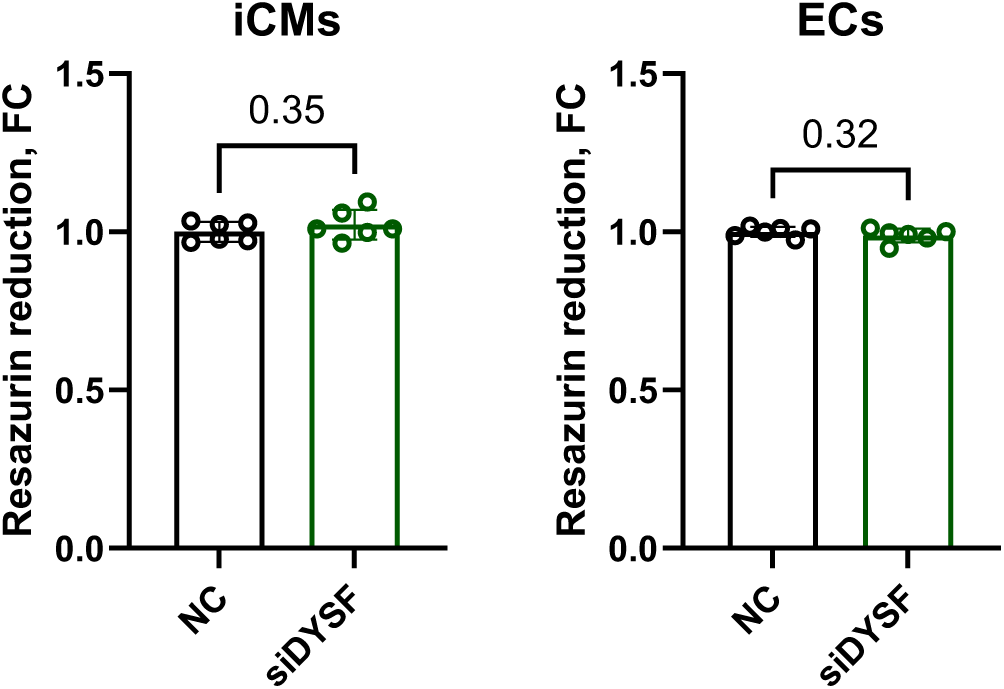
Effect of *DYSF* silencing on resazurin reduction in induced cardiomyocytes (iCMs) and endothelial cells (ECs) 72 h after transfection (n=6; unpaired t-test; mean ± SD).

**Supplementary Figure 7.**
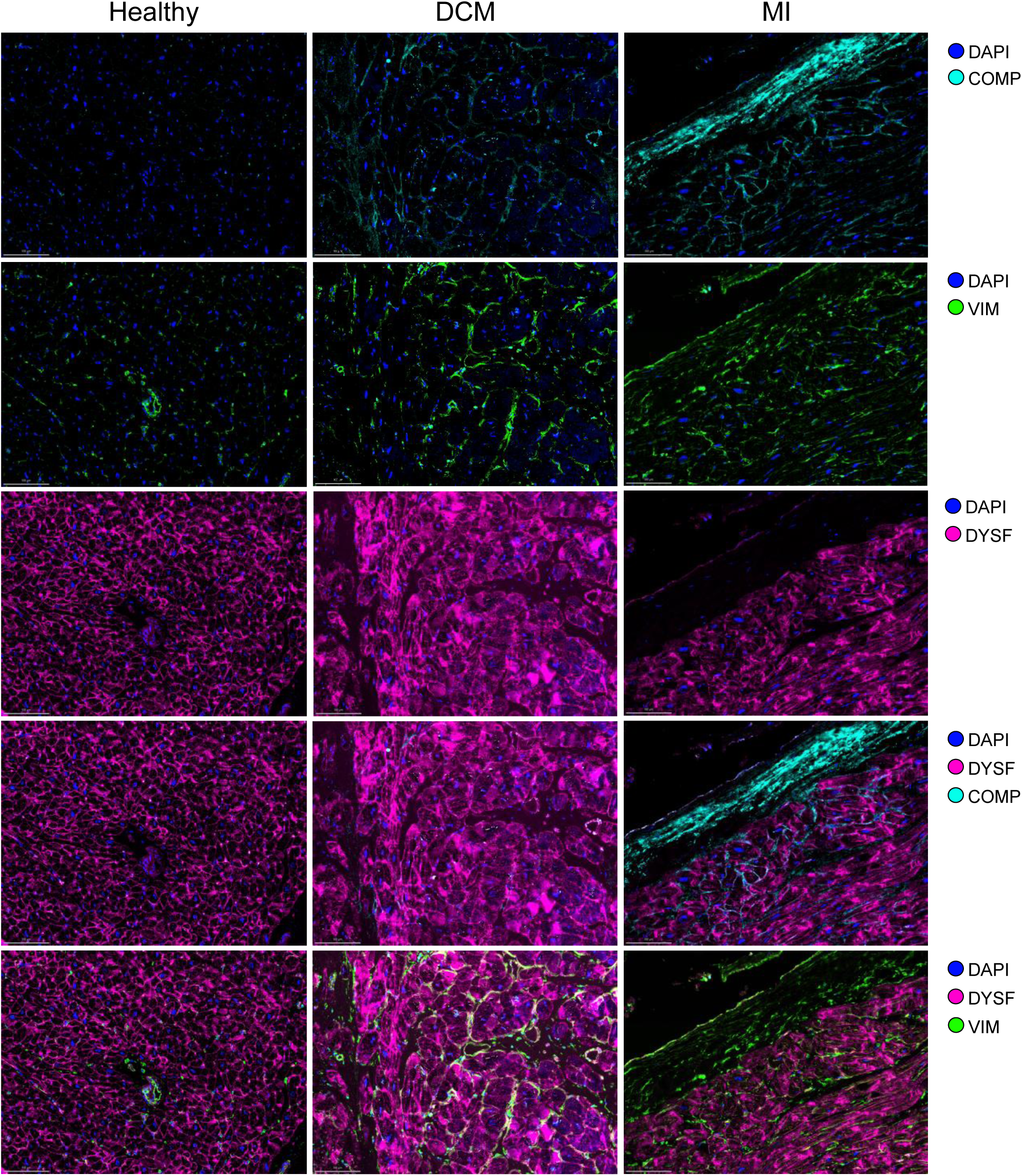
Multiplex immunofluorescence of human myocardium. Formalin-fixed, paraffin-embedded sections of DCM, MI and healthy human myocardium were stained using the COMET multiplex immunofluorescence platform (Lunaphore). Representative images are shown. Scale bar: 100 µm.

**Supplementary Figure 8.**
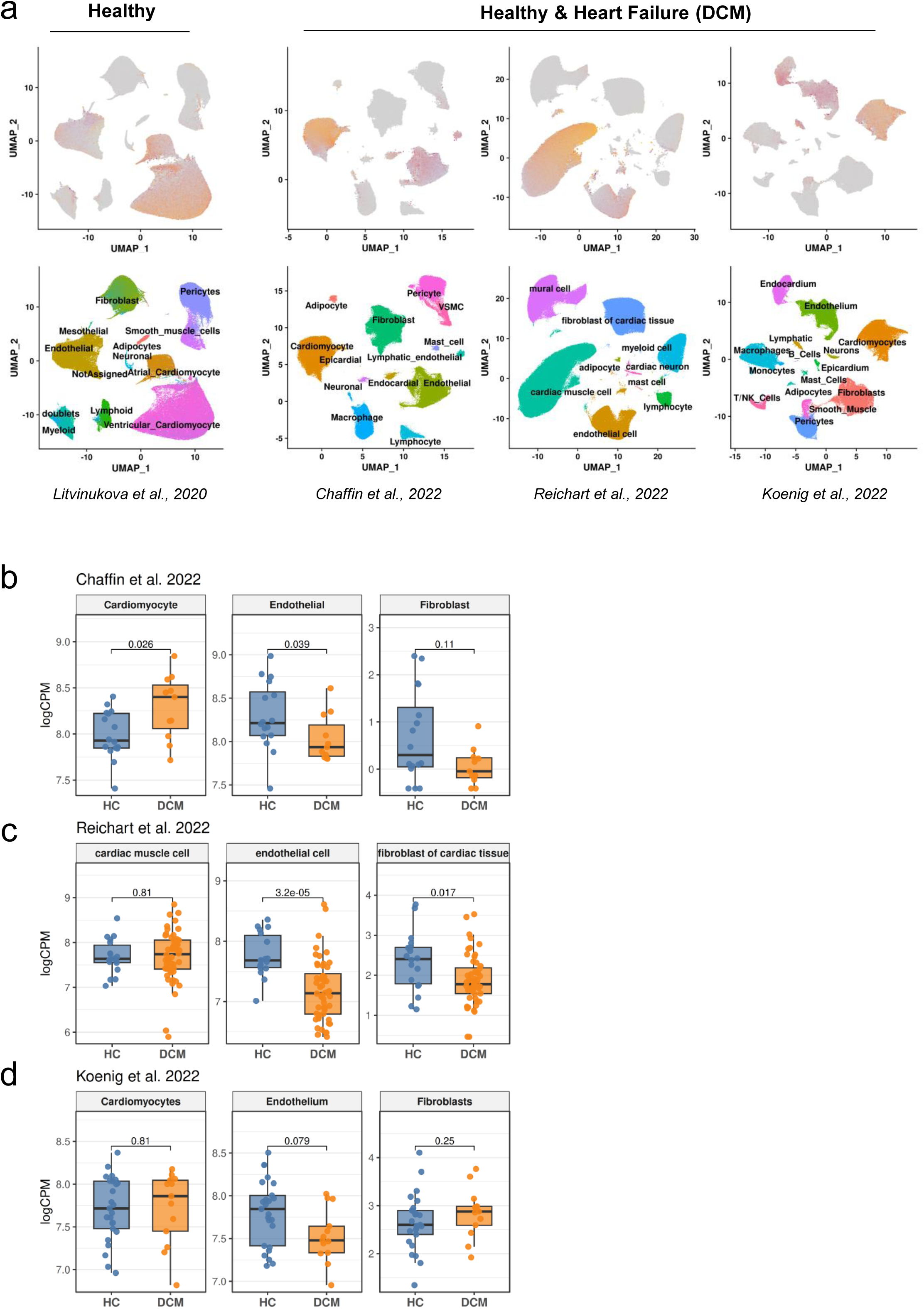
*DYSF* expression in human healthy and HF myocardium across publicly available transcriptomic datasets**. (a)** *DYSF* expression in myocardial cell types. **(b-d)** Differential expression of *DYSF* in HF in Chaffin et al. (SCP1303, **b**), Reichart et al. (EGAS00001006374, **c**), and Koenig et al. (GSE183852, **d**) datasets.

**Supplementary Figure 9.**
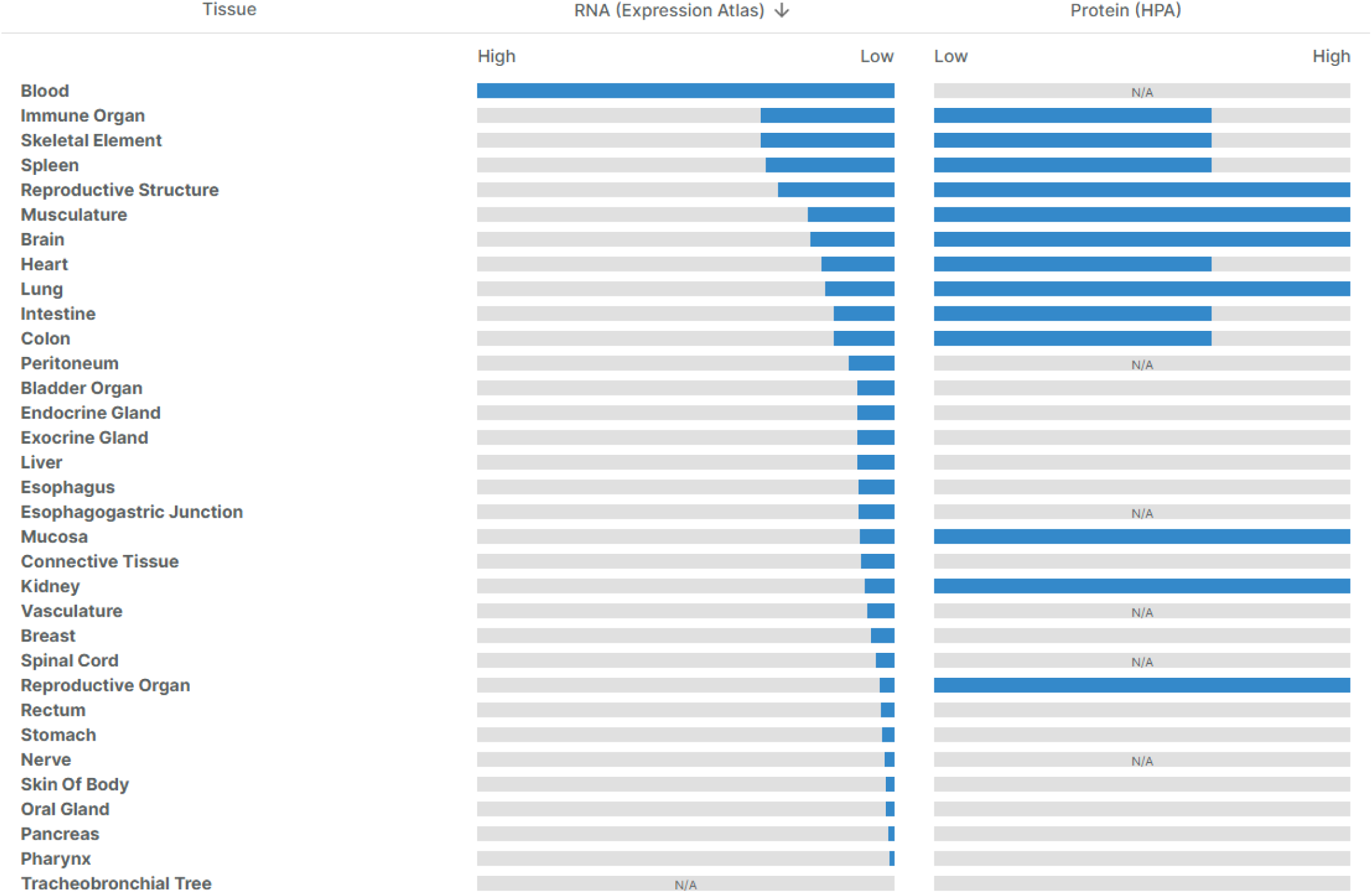
Baseline RNA expression and protein levels for dysferlin. Source: ExpressionAtlas, HPA and GTEx. Retreived from the Open Target Platform (https://platform.opentargets.org/target/ENSG00000135636, accessed on 10.11.2025).

**Supplementary Figure 10.**
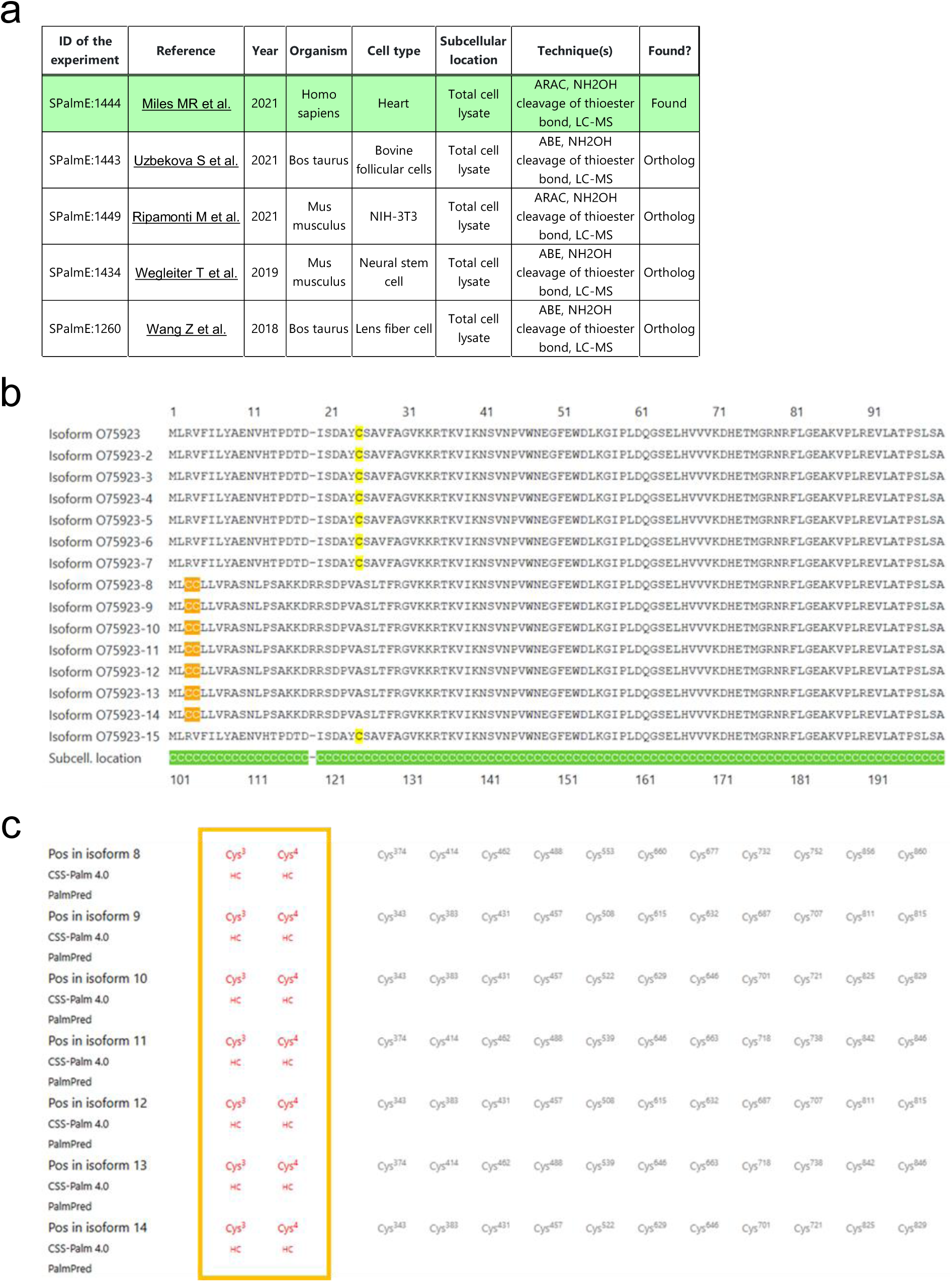
Dysferlin is a putative S-acylated protein. **(a)** Top 5 palmitoyl-proteome experiments predicting dysferlin to be S-acylated, according to SwissPalm database. **(b)** Predicted S-acylation sites in the dysferlin protein (O75923, DYSF_HUMAN). Partial dysferlin isoform sequence alignment with cysteines highlighted in yellow (standard) or orange (predicted S-acylation sites). Green-highlighted ‘C’ indicates cytosolic/nuclear localization. **(c)** Cysteine summary and predictions showing high-confidence (HC, red font) cysteine residues. Data retrieved from the SwissPalm database. Source: https://swisspalm.org/proteins/O75923, accessed on 20.10.2024. ARAC: acyl-resin-assisted capture, ABE: acyl-biotin exchange assay, NH2OH: hydroxylamine.

